# Bromodomain Factor 5 as a Target for Antileishmanial Drug Discovery

**DOI:** 10.1101/2023.08.21.554125

**Authors:** Catherine N. Russell, Jennifer L. Carter, Juliet M. Borgia, Jacob Bush, Félix Calderón, Raquel Gabarró, Stuart J. Conway, Jeremy C. Mottram, Anthony J. Wilkinson, Nathaniel G. Jones

**Affiliations:** York Structural Biology Laboratory and York Biomedical Research Institute, Department of Chemistry, University of York, York, YO10 5DD, UK; Department of Chemistry, Chemistry Research Laboratory, University of Oxford, Mansfield Road, Oxford, OX1 3TA, UK; GSK, Gunnels Wood Road, Stevenage, Hertfordshire, SG1 2NY, UK; The Francis Crick Institute, London, NW1 1AT, UK; GSK Global Health, Tres Cantos, 28760 Madrid, Spain; York Biomedical Research Institute, Department of Biology, University of York, York, YO10 5NG, UK

**Keywords:** Leishmania, bromodomain, epigenetics, drug discovery, fluorescence polarisation

## Abstract

Leishmaniases are a collection of neglected tropical diseases caused by kinetoplastid parasites in the genus *Leishmania*. Current chemotherapies are severely limited and the need for new antileishmanials is of pressing international importance. Bromodomains are epigenetic reader domains that have shown promising therapeutic potential for cancer therapy and may also present an attractive target to treat parasitic diseases. Here, we investigate *Leishmania donovani* bromodomain factor 5 (*Ld*BDF5) as a target for antileishmanial drug discovery. *Ld*BDF5 contains pair of bromodomains (BD5.1 and BD5.2) in an N-terminal tandem repeat. We purified recombinant bromodomains of *L. donovani* BDF5 and determined the structure of BD5.2 by X-ray crystallography. Using a histone peptide microarray and fluorescence polarisation assay, we identified binding interactions of *Ld*BDF5 bromodomains with acetylated peptides derived from histones H2B and H4. In orthogonal biophysical assays including thermal shift assays, fluorescence polarisation and NMR, we showed that BDF5 bromodomains bind to human bromodomain inhibitors SGC-CBP30, bromosporine and I-BRD9, moreover, SGC-CBP30 exhibited activity against *Leishmania* promastigotes in cell viability assays. These findings exemplify the potential BDF5 holds as a drug target in *Leishmania* and provide a foundation for the future development of optimised antileishmanial compounds targeting this epigenetic reader protein.

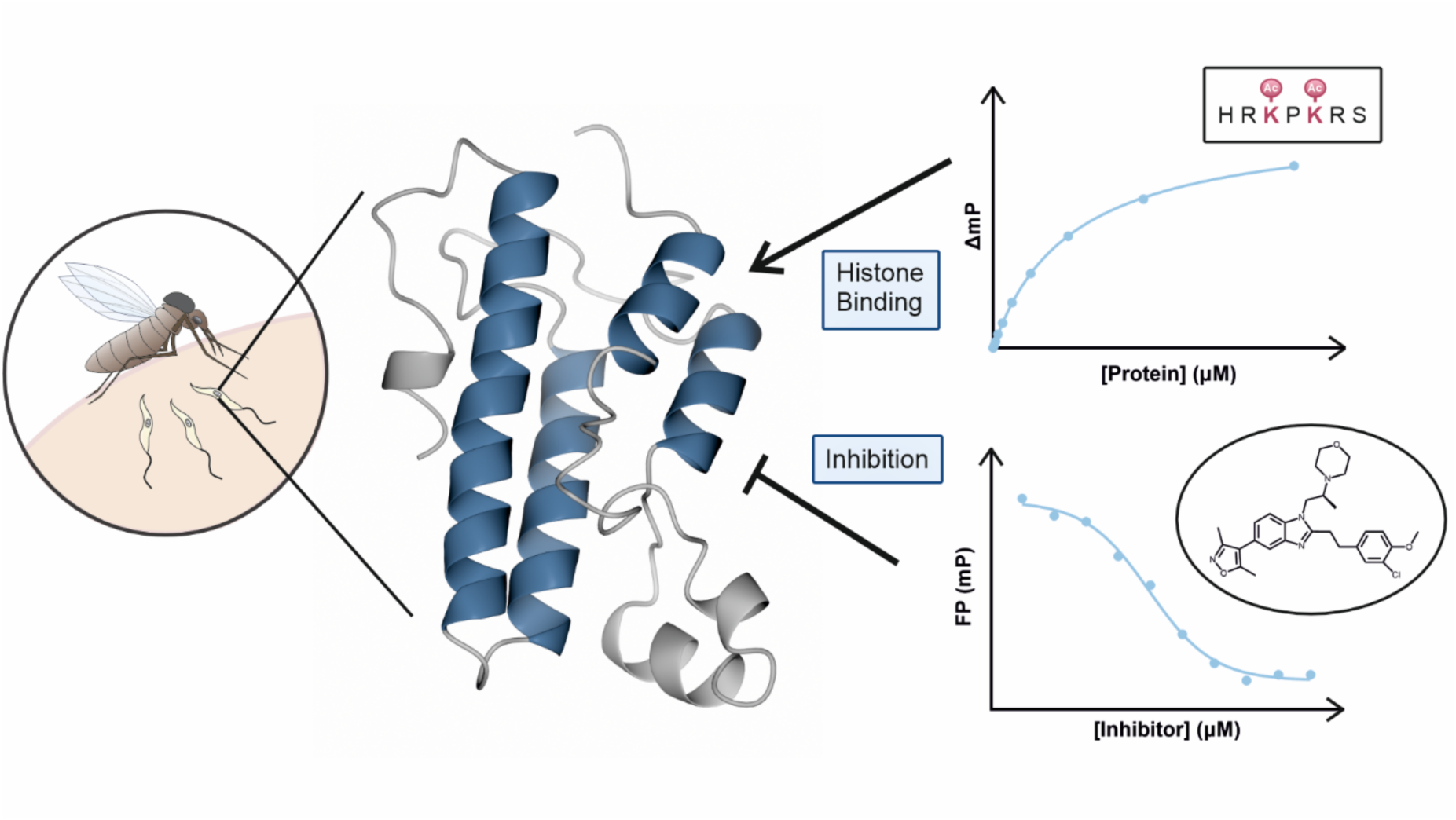

## Introduction

Leishmaniasis is a neglected tropical disease which is endemic in approximately 100 countries and had an estimated prevalence of over four million cases in 2019^1,2^. The three most prevalent forms of leishmaniasis are cutaneous, mucocutaneous and visceral. Visceral leishmaniasis (VL) is the most severe form of the disease, in which the parasite infects organs including the spleen and liver, typically causing anaemia or intercurrent bacterial infection which can be fatal if untreated. VL is predominantly caused by *Leishmania donovani* and *L. infantum* species, 500,000 new cases of VL and 50,000 deaths are estimated to occur annually^1,3^. Current treatments for leishmaniasis include pentavalent antimonials, amphotericin B, miltefosine, paromomycin, and pentamidine, with all but miltefosine requiring parenteral administration. These chemotherapies suffer from high costs, long treatment times, toxicity, and growing resistance^4^. Ongoing public-private partnerships have been successful in advancing new chemical entity antileishmanials into clinical trials – all of which were identified by phenotypic screening and target deconvolution^5,6^. However, owing to various challenges, target-based antileishmanial drug discovery has shown limited success, and despite extensive research, the target proteins of most antileishmanials currently remain unknown, primarily due to a lack of genetically validated targets to feed into drug discovery programs^7,8^.

*Leishmania* parasites are transmitted by phlebotomine sand flies and follow a complex lifecycle involving differentiation between amastigote and promastigote forms. Promastigotes are the motile form found in the insect vector, while the amastigote form is the clinically relevant intracellular form found in the mammalian host^9^. Parasite survival and infectivity is highly dependent on the ability of the parasite to transition between these structurally and phenotypically distinct stages. This is achieved by enacting precise control of gene expression. Epigenetic transcriptional regulation is a mechanism in which gene expression is regulated by modification of DNA or post-translational modifications (PTMs) of histone proteins in nucleosomes. One such PTM is the acetylation of lysine residues on histone tails, typically associated with an open chromatin structure and active gene transcription. Lysine acetylation is catalysed by histone acetyltransferases (HATs) and deacetylation by histone deacetylases (HDACs), while the ‘reader’ proteins of acetylated lysine (acetyl-lysine) commonly feature a module known as the bromodomain^10,11^. Because of their polycistronic genome arrangement, *Leishmania* exhibit limited differential transcriptional regulation to achieve distinct gene expression states, mostly relying on post-transcriptional processes and genome plasticity^12,13^. Therefore, the regulation of polymerase II activity by epigenetic processes is mostly likely to be global^14,15^, and disruption of this process would likely be lethal for all transcriptionally active stages of the parasite.

Bromodomains contain ~ 110 amino acid residues which fold to form a bundle of four antiparallel α-helices (αZ, αA, αB and αC), connected in a left-handed topology. Two variable loops (ZA and BC) form a hydrophobic acetyl-lysine binding pocket containing conserved asparagine and tyrosine residues involved in recognition of the acetyl-lysine substrate. The binding pocket also often contains a network of water molecules implicated in binding substrates^16,17^. Binding specificity is determined by sequence variations in the ZA and BC loops, and whilst affinity for a single acetyl-lysine is low, flanking residues in the histone substrate provide additional specificity determinants and contribute to a higher affinity interaction^18–21^.

Bromodomains, particularly the bromodomain and extra-terminal (BET) family, have been comprehensively studied in humans where they play well-documented roles in chromatin remodelling and regulation of gene expression^22^. Bromodomain proteins are important for cellular homeostasis, and enact their regulatory functions through a range of mechanisms, including acting as scaffolds for the recruitment of other proteins to DNA or acting as transcription co-modulators. Additionally, bromodomains are often found in multidomain proteins alongside catalytic domains such as acetyltransferase and methyltransferase domains^19,23^. Dysregulation of bromodomains has been associated with a myriad of diseases including cancers, such as leukaemia, NUT midline carcinoma, and breast cancer^24^. As a result, intensive research efforts have been directed towards the discovery and development of bromodomain inhibitors as anti-cancer drugs^16,25^. These efforts have identified potent and selective bromodomain-targeting molecules, many of which have progressed into clinical trials^26,27^. These successes exemplify the tractability of bromodomains as ligandable drug targets, and indicate that these reader domains also offer an avenue for the development of antileishmanials.

Five canonical bromodomain factor (BDF) proteins are encoded in the *Leishmania* genome (BDF1-5) alongside an additional three non-canonical bromodomains (BDF6-8). The conserved asparagine and tyrosine residues are identifiable in the five canonical proteins, and each contains a single bromodomain with the exception of BDF5, which contains tandem bromodomains (BD5.1 and BD5.2). BDF5 is also predicted to contain a C-terminal MRG (MORF4-related gene) domain. Elsewhere, these domains function in chromatin remodelling and transcription regulation proteins^28^. Recently, we found that BDF1-5 are essential for *L. mexicana* promastigote survival, as these genes were refractory to Cas9-targeted gene deletion. Furthermore, inducible knockout using a dimerisable split Cre recombinase (DiCre) system showed that BDF5 is essential for both promastigote survival and murine infection competence. BDF5, which is expressed in both *Leishmania* lifecycle stages, localises to the nucleus where it has an essential role in RNA polymerase II-mediated transcription. The protein is enriched at transcriptional start regions (TSRs), with RNA-seq analysis revealing a global decrease in transcription following BDF5 deletion. Using an *in situ* proximity labelling technique (XL-BioID), the proximal proteome of BDF5 was found to include 156 proteins with roles that include epigenetic regulation of transcription, mRNA maturation, and DNA damage repair. Four other BDF proteins appeared in the proximal proteome, of which BDF3, BDF6, and BDF8 were validated by coimmunoprecipitation^14^. Based on these findings, and related data from immunoprecipitation experiments in *T. brucei*^29^, it was proposed that BDF5 is a component of a protein assembly that includes BDF3, BDF8, and HAT2. This Conserved Regulators of Kinetoplastid Transcription (CRKT) complex associates with TSRs and influences transcription^14^.

Research has already begun to shed light on bromodomains in other parasitic protozoa^30^. BDF orthologues have been identified in *Trypanosoma cruzi* and *T. brucei*, where their inhibition using small molecule ligands has been investigated. Recombinant bromodomain from *T. cruzi* BDF3 interacts with the human bromodomain inhibitors JQ1 and I-BET151, while *T. brucei* BDF2 recombinant bromodomain binds I-BET151 and GSK2801, with exposure to these compounds resulting in disrupted parasite growth and abnormal lifecycle progression^31–33^. GSK2801 was also shown to prevent binding of BDF2 to its acetylated histone substrate^31^. Novel ligands have also been investigated, for example, fragment-based approaches were applied to identify tool compounds which bind to *T. cruzi* BDF3^34^. These discoveries, in conjunction with the recent genetic target validation of *Leishmania* BDF5, and the traction bromodomain inhibitors have gained in cancer research, provide a strong rationale for the investigation of BDF5 as a drug target in *Leishmania*.

Here, we report the interactions of recombinant bromodomains BD5.1 and BD5.2 of BDF5 with acetylated histone-derived peptides and small molecule inhibitors. We identified acetylated sequences in *Ld*H2B and *Ld*H4 which interact with the protein. Furthermore, we investigated the binding of BD5.1 and BD5.2 to human bromodomain inhibitors, including bromosporine, SGC-CBP30, and I-BRD9, in orthogonal biophysical assays. In addition, we determined the crystal structure of unliganded (apo) BD5.2, enabling comparison with the structure of a BD5.2-bromosporine complex. Promisingly, we show that that the compound SGC-CBP30 not only binds to BD5.1, but that it also elicits an inhibitory effect on *Leishmania* promastigotes in cell viability assays. Using these approaches, we demonstrate the potential of BDF5 as a target for the development of new antileishmanial compounds.

## Results & Discussion

### LdBDF5 Recombinant Protein Production and Structural Characterisation

We first sought to generate soluble recombinant protein for the bromodomains of *L. donovani* BDF5 (LDBPK_091320); BD5.1 containing the first bromodomain, BD5.2 containing the second bromodomain, and BD5T containing the tandem bromodomain pair (Figure 1A; Table S1–3). Plasmids derived from pET-15-MHL containing the bromodomain coding sequences were used to direct IPTG-induced overproduction of the proteins in *E. coli*. The recombinant proteins were purified using immobilized metal affinity chromatography (IMAC) and size exclusion column chromatography (Figure 1B). Size exclusion chromatography with multi-angle laser light scattering (SEC-MALLS) analysis revealed that all three proteins were monomeric with molecular masses consistent with expected values (Figure 1C). The bromodomain of *L. donovani* BDF2 (LDBPK_363130), herein referred to as BD2, was also recombinantly produced and analysed by the same methods (Table S4; Figure S1).

**Figure 1.**
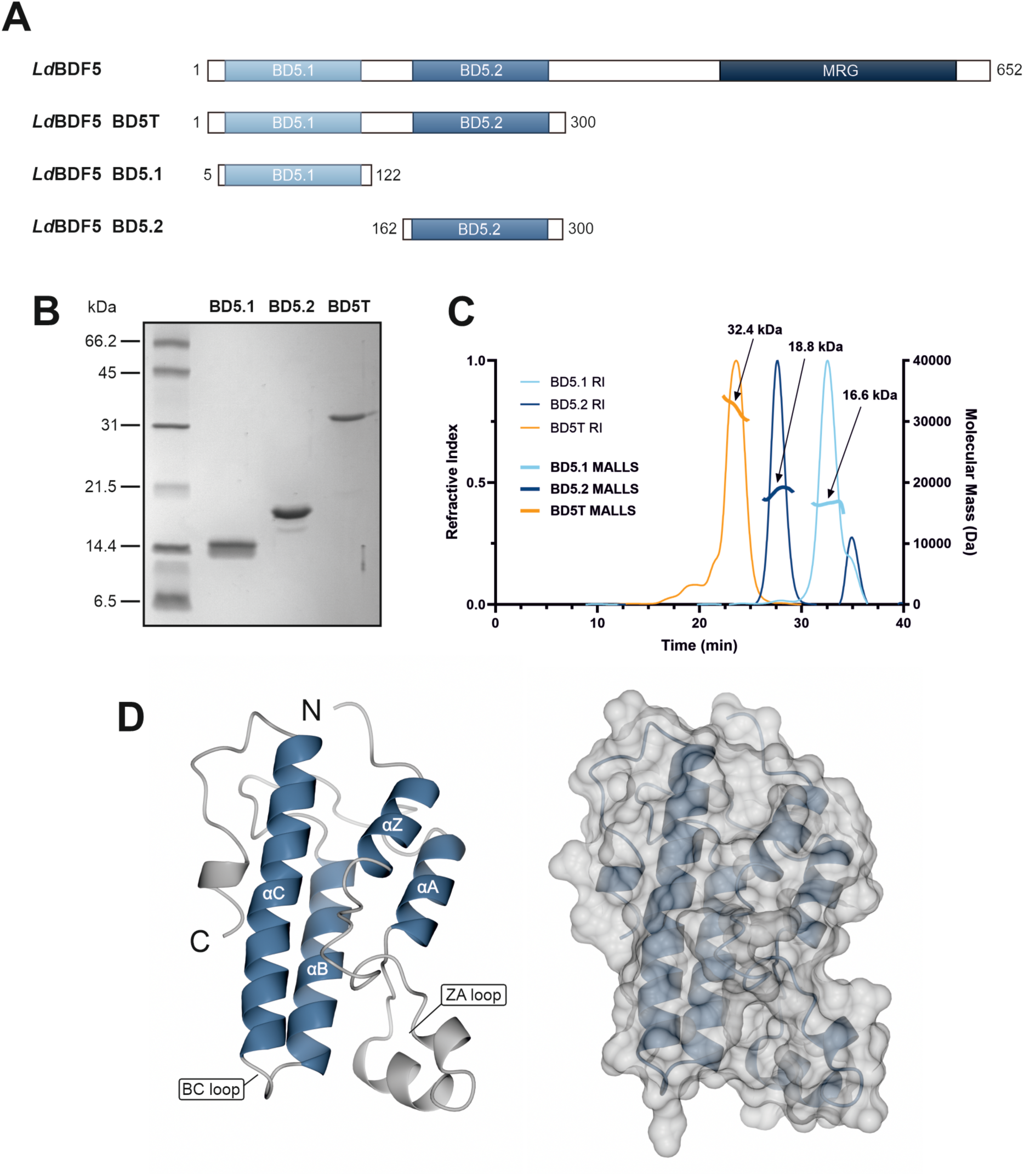
(A) Domain architecture of native *L. donovani* BDF5 and bromodomains in the three recombinant protein constructs (BD5T, BD5.1 and BD5.2), where MRG indicates a predicted C-terminal MORF4-related gene C-terminal domain. (B) 17.5% SDS-PAGE analysis of purified recombinant proteins BD5.1, BD5.2 and BD5T, with expected molecular mass of 15.2, 17.9 and 33.0 kDa, respectively. (C) SEC-MALLS analysis of recombinant *Ld*BDF5 proteins; BD5.1 (light blue), BD5.2 (dark blue) and BD5T (orange), showing the refractive index (RI) with arrows indicating MALLS curves labelled with the associated estimated molecular masses (in all cases within 10% of expected values). The chromatogram shows elution time on the x axis, refractive index (RI) on the left y axis and molecular mass on the right x axis. (D) X-ray structure of BD5.2 (PDB code 8BPT) displaying ribbon and surface representations; figures generated using CCP4mg software^40^.

**Table 1.** *K*_d_ values for binding of H2B and H4 peptides to BD5.1 and BD5.2 determined by FP. Binding affinities calculated by plotting protein concentration against mean, blank-corrected ΔmP values and fitting one-site specific binding non-linear regression models. *K*_d_ values are reported ± standard error.

| Peptide | BD5.1 $K_d$ (μM) | BD5.2 $K_d$ (μM) |
| --- | --- | --- |
| H2B<sub>9-23</sub>K9<sup>Ac</sup>K15<sup>Ac</sup>K19<sup>Ac</sup>K21<sup>Ac</sup> | 117 ± 2.3 | 463 ± 45 |
| H2B<sub>8-16</sub>K9<sup>Ac</sup>K15<sup>Ac</sup> | 170 ± 10 | 366 ± 60 |
| H2B<sub>17-23</sub>K19<sup>Ac</sup>K21<sup>Ac</sup> | 77.5 ± 0.95 | 160 ± 10 |
| H4<sub>1-15</sub>K2<sup>Ac</sup>4<sup>Ac</sup>10<sup>Ac</sup>14<sup>Ac</sup> | 137 ± 4.4 | 353 ± 49 |

The high-quality recombinant protein allowed us to determine the structure of BD5.2 by X-ray crystallography (Figure 1D; Table S5; PDB code 8BPT). The two (A and B) chains of BD5.2 in the asymmetric unit can be overlaid using SSM superpose routines to give an rmsΔ of 0.98 Å for 139 equivalent atoms. Each chain adopts the canonical bromodomain fold, comprising four anti-parallel alpha helices with a prominent ZA loop (joining helices αZ and αA) containing additional helical elements. Comparison of this structure with that of BD5.2 bound to bromosporine (PDB code 5TCK; Figure S2C) shows that few conformational changes take place upon inhibitor binding; overlay of chains A from the two structures by SSM superposition gives an rmsΔ of 0.52 Å for 139 matched atoms. BD5.2 is the second bromodomain of the *Ld*BDF5 tandem bromodomain pair, and also exhibits high sequence and structural similarity with the first bromodomain, BD5.1; overlay of chain A in the unliganded BD5.2 structure with chain A in a BD5.1 co-crystal structure (in complex with SGC-CBP30; PDB code 6BYA) by SSM superposition gives an rmsΔ of 1.19 Å for 112 matched atoms.

As no full-length *Leishmania* BDF structure has been determined experimentally, we used AlphaFold to predict the structure of *Ld*BDF5^35,36^ (Figure S3). The proposed structure has the two bromodomains positioned next to each other, with the binding pockets facing outwards at a slight angle to one another. This is similar to the arrangement seen in X-ray structures of tandem bromodomains of human Rsc4 and TAF1^37,38^. The other region in the AlphaFold model predicted with confidence is the MRG domain, which potentially functions in chromatin remodelling and transcription regulation^28^. Recent work also appears to demonstrate that the MRG domain is capable of mediating CRKT protein interactions in trypanosomatids^39^. Whilst the conserved folds of the two bromodomains and the MRG domain are predicted with confidence (pLDDT > 70), the structures of segments of the polypeptide linking the domains are not. As a result, the spatial orientation of the three domains is only tentatively assigned.

### Identifying Histone Binding Interactions of LdBDF5

The *Ld*BDF bromodomains are predicted to bind acetyl-lysine residues on histone tails, however, their specific targets have not been established. Here, we explored their potential to engage acetylated histone peptides experimentally in an unbiased manner, using an array that could be probed with recombinant bromodomains. As kinetoplastid histone tails are significantly divergent from those of mammalian orthologues^41,42^, a custom peptide microarray was developed. In the absence of a comprehensive dataset of *Leishmania* histone PTMs, the microarray represented a panel of around 1000 unmodified, pan-acetylated and mono-acetylated peptides derived from the 25 N-terminal residues of *L. donovani* histones, tiled in 15-mer duplicates to cover the full sequence. These were probed with recombinant BD5.1, BD5.2, and BD2 (Figure 2A).

**Figure 2.**
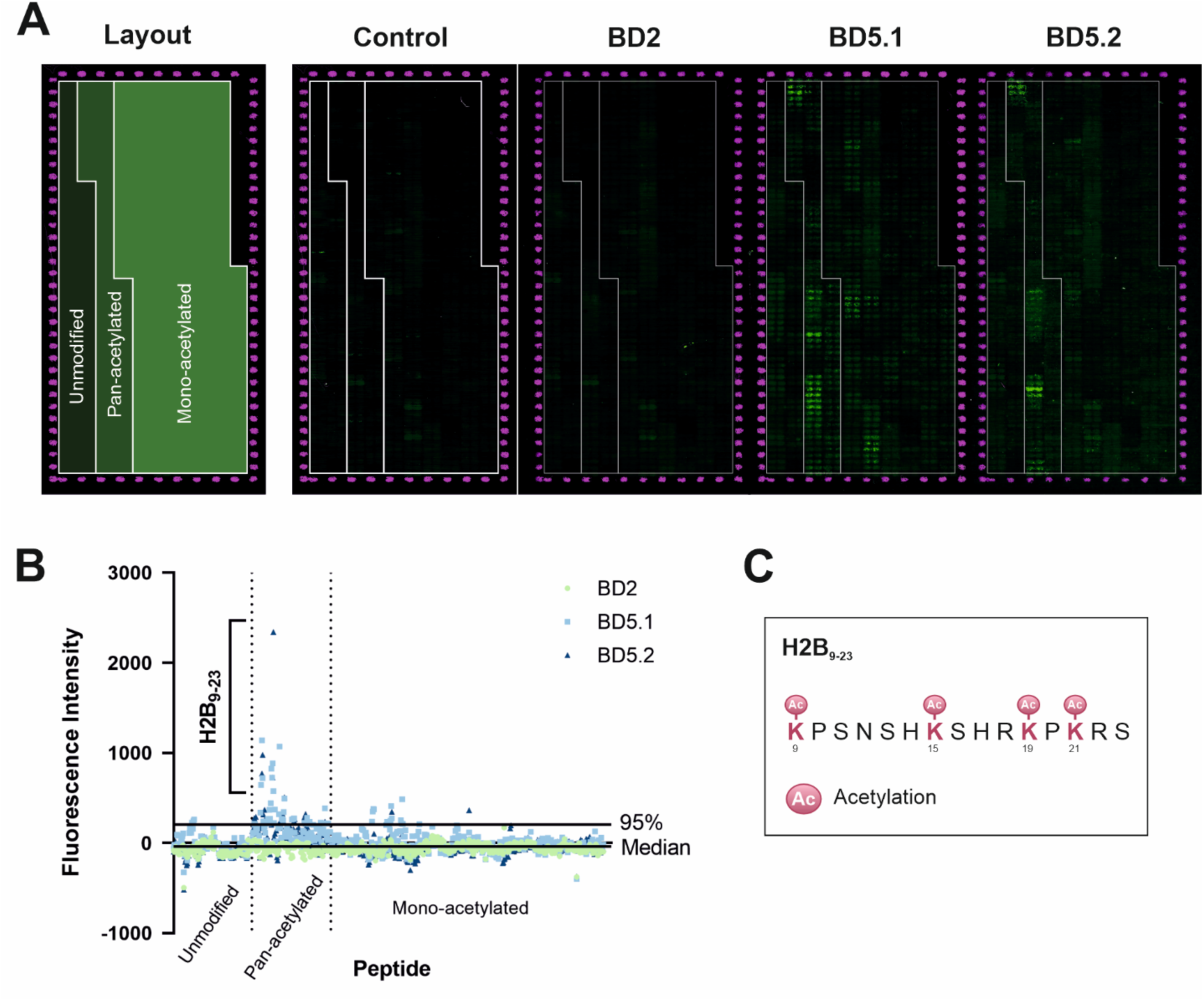
(A) Image of histone peptide microarray output from microarray scanner; glass slide containing microarrays synthesised by PepperPrint and probed with recombinant *L. donovani* bromodomain proteins followed by anti-6xHis (green) and anti-HA (red) fluorescent conjugated antibodies. The control array was probed with the antibodies alone to exclude non-specific binding. (B) Blank-corrected fluorescence intensity readings from the histone peptide microarray plotted against peptide number for BD2, BD5.1 and BD5.2 with median and 95th percentile indicated. A pan-acetylated H2B sequence interacting with BD5.1 and BD5.2 is evident. (C) Sequence of the pan-acetylated peptide based on the H2B_9-23_ sequence.

Plotting the mean fluorescence signal from duplicate peptide spots against peptide number identified a pan-acetylated 15 amino acid residue sequence in the N-terminal tail of histone H2B, which appeared to bind both BD5.1 and BD5.2 (Figure 2B), with signal intensities in the top 5% of the distribution. In comparison, no signal was detected for the same peptide with BD2 or in the antibody-only control, suggesting that these BD5-peptide interactions are specific. This H2B_9-23_ sequence contained four acetyl-lysines; K9, K15, K19 and K21. In *T. brucei*, minor acetylation has been detected at H2BK12 and K16 which correspond to K15 and K19 in H2B from *L. donovani*^43^. In a thermal shift assay (TSA), a pan-acetylated peptide based on this sequence (Figure 2C) gave rise to a small increase in the melting temperature of BD5T (Δ*T*_m_ = 0.60 ± 0.15 °C; unpaired t-test, P < 0.001, n = 6) (Figure S4A) when tested at 400 µM, indicative of ligand binding and an associated increase in protein stability. In comparison, an unmodified version of the peptide produced a markedly smaller thermal shift under the same conditions (Δ*T*_m_ = 0.19 ± 0.09 °C; unpaired t-test, P = 0.0232, n = 6) (Figure S4B). This tends to corroborate the peptide microarray data and prompted further exploration of an interaction between *Ld*BDF5 and H2B.

### Characterising Interactions of LdBDF5 with Histones H2B and H4

The *in vivo* acetylation status of the identified H2B sequence is unknown, however, HAT acetylation sites have been identified in *L. donovani* histone H4. Both HAT1 and HAT2 have been found to acetylate H4K10^44,45^, while H4K4 is an acetylation site of HAT2 and HAT3^46,47^. Additionally, H4K14 was identified as a major acetylation site and H4K2 as a potential minor acetylation site of HAT4^48^. Peptides from histone H4 were therefore investigated alongside those from histone H2B in the search for BDF5 binding sequences.

To characterise interactions of *Ld*BDF5 with H2B and H4, fluorescently-labelled peptides derived from the N-terminal regions of these histones were designed and synthesised for application in a fluorescence polarisation (FP) assay (Figure 3A). These included the tetra-acetylated 15mer H2B sequence identified in the microarray and a tetra-acetylated 15mer region of H4 containing the four lysines previously identified as HAT acetylation sites. Additional, shorter H2B peptides were also synthesised each containing two acetyl-lysines. Unmodified peptides were also included for the purpose of establishing binding specificity. FP protein-binding experiments were performed in which a fixed concentration of peptide (800 nM) was titrated against increasing concentrations of recombinant *Ld*BDF proteins up to 300 µM, with binding measured as an increase in FP, and associated *K*_d_ values calculated (Figure 3B-E; Table 1).

**Figure 3.**
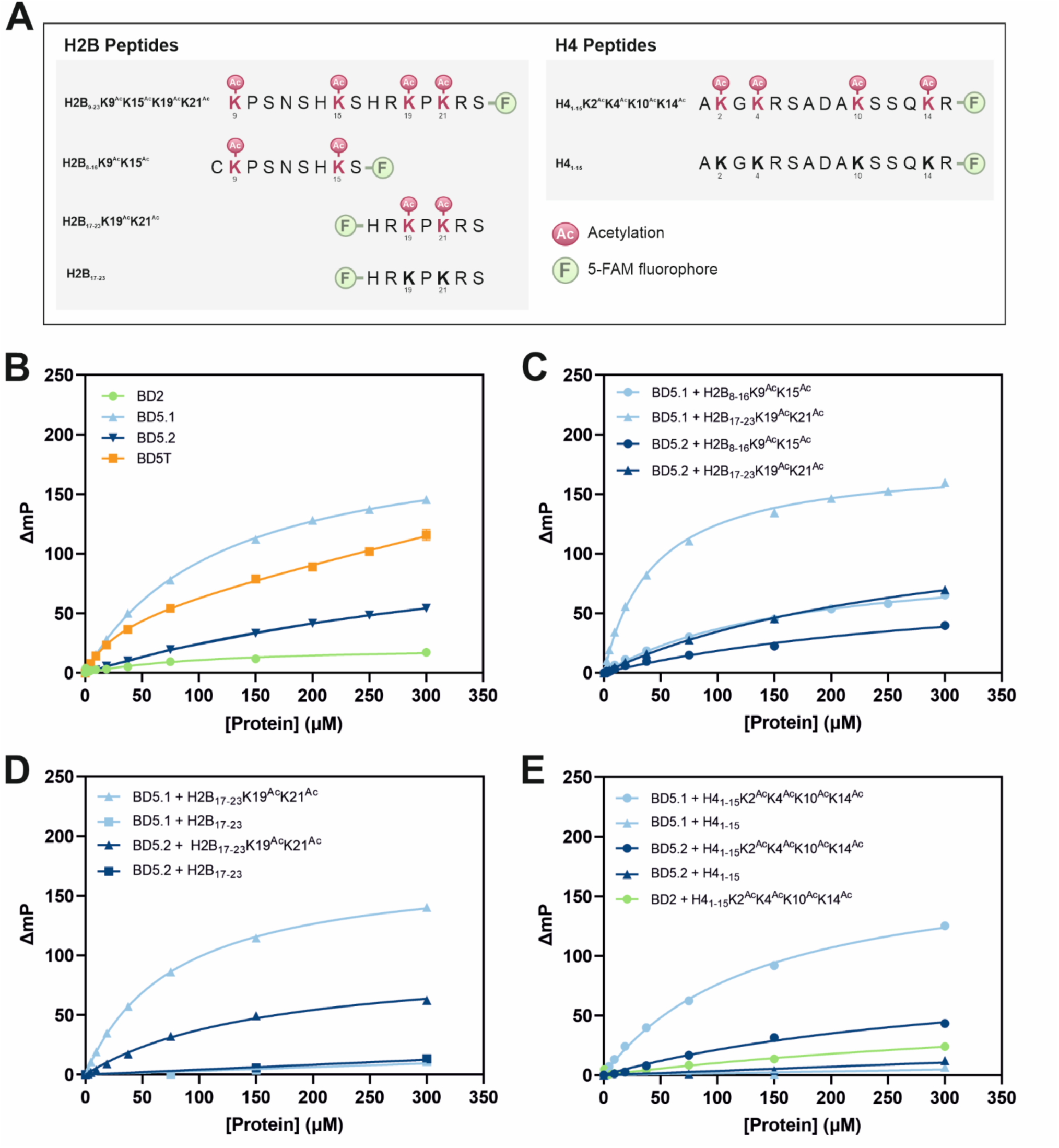
(A) Sequences of peptides derived from *L. donovani* histones H2B and H4. F denotes a conjugated 5-carboxyfluorescein (5-FAM) fluorophore. Ac indicates acetylation of a lysine residue. FP protein-peptide binding curves for (B) H2B_9-23_K9^Ac^K15^Ac^K19^Ac^K21^Ac^ with BD2, BD5.1, BD5.2 and BD5T; (C) H2B_8-16_K9^Ac^K15^Ac^ and H2B_17-23_K19^Ac^K21^Ac^ with BD5.1 and BD5.2; (D) H2B_17-23_K19^Ac^K21^Ac^ and unmodified H2B_17-23_ with BD5.1 and BD5.2; and (E) H4_1-15_K2^Ac^4^Ac^10^Ac^14^Ac^ with BD5.1, BD5.2 and BD2, and unmodified H4_1-15_ with BD5.1 and BD5.2. In B-E protein concentration is plotted against mean, blank-corrected ΔmP values, fitted to one- or two-sites specific binding non-linear regression curves for single or tandem bromodomain proteins respectively. Error bars represent SD (n = 3); these are largely invisible on account of low SD values.

The *Ld*BDF5-H2B association was first investigated using the 15mer **H2B_9-23_K9^Ac^K15^Ac^K19^Ac^K21^Ac^** peptide. An FP response was observed with BD5.1, BD5.2 and BD5T, but not with BD2 (Figure 3B), consistent with the peptide microarray data. The dissociation constant calculated for BD5.1 was approximately 4-fold higher than for BD5.2 (Table 1). The 15mer sequence was subsequently split into two di-acetylated peptides, **H2B_8-16_K9^Ac^K15^Ac^** and **H2B_17-23_K19^Ac^K21^Ac^**, assayed for binding to BD5.1 and BD5.2 (Figure 3C). FP binding curves showed that the **H2B_17-23_K19^Ac^K21^Ac^** peptide exhibited the strongest binding to both bromodomains. Finally, to confirm selectivity of binding to the H2B 7mer sequence, the acetylated peptide, **H2B_17-23_K19^Ac^K21^Ac^**, was tested alongside an unmodified version, **H2B_17-23_** (Figure 3D; Table 1). As seen in the previous FP experiment, the acetylated peptide bound to both BD5.1 (*K*_d_ = 77.5 ± 0.95 µM) and BD5.2 (*K*_d_ = 160 ± 10 µM), whereas no significant FP response was elicited by the unmodified peptide, indicating that binding is modification-specific. Regarding a potential binding mechanism of *Ld*BDF5 with **H2B_17-23_K19^Ac^K21^Ac^**, the presence of multiple acetyllysines in this sequence could indicate a cooperative binding mechanism with simultaneous engagement of both bromodomains. This peptide exhibited stronger binding to BD5.1 compared with BD5.2, which led us to believe that the interaction primarily involves binding of the first bromodomain to one or both of the acetylated K19 and K21 residues, with BD5.2 potentially facilitating binding via an additional weaker association with the histone.

The H4-derived 15mer peptides were then analysed for binding to BD5.1, BD5.2 and BD2 using the same approach (Figure 3E). The acetylated peptide **H4_1-15_K2^Ac^K4^Ac^K10^Ac^K14^Ac^** displayed binding to BD5.1 (*K*_d_ = 137 ± 4.4 µM). By contrast, the same peptide produced a very minor FP response with BD5.2 and an almost negligible response with BD2. The unmodified peptide **H4_1-15_** did not bind to BD5.1 or BD5.2, again indicating that the interactions are specific. From these findings, it may be inferred that in addition to H2B, *Ld*BDF5 also has the capacity associate with H4 in an interaction predicted to involve the first bromodomain and acetylated K2, K4, K10 and/or K14 in the histone.

Although the status of H2B acetylation in *Leishmania* is yet to be determined, it has been reported that H2B can be acetylated in *T. brucei* at K12 and K16^43^. Multiple N-terminal H2B acetylation sites have also been reported and shown to mark active promoters in humans^49^ and also murine enhancers^50^. Although further work is required to verify, it may be the case that acetylated H2B and H4 at transcriptional start regions in *Leishmania* can act to recruit BDF5, which is consistent with the known ability of BDF5 to be associated with the broad transcriptional start regions *of L. mexicana* and promote gene expression.

### Targeting LdBDF5 with Human Bromodomain Inhibitors

In addition to exploring the histone binding interactions of *Leishmania* BDF5 bromodomains, we also sought to establish whether the protein is a viable target for the development of antileishmanial compounds. The recent surge in bromodomain research has led to the availability of numerous compounds suitable for use as chemical probes^25,51^. These include many human bromodomain inhibitors which have been applied to probe the biological functions of bromodomains in other organisms, including parasites^52^. Our aim was to utilise these existing tool compounds and apply them within biophysical assays for the identification of small-molecule inhibitors of BDF5 to support drug discovery.

We screened a panel of 15 commercially available bromodomain inhibitors (Figure S5), against BD5.1, BD5.2 and BD5T using TSA, a technique which has also been used to identify ligands of human bromodomains^53^. Included in the panel were compounds targeting bromodomains in all eight human bromodomain families with up to nanomolar binding affinities (*K*_d_), ranging in molecular mass from 347 Da (PFI-1) to 527 Da (GSK8814). Notable inclusions were the pan-bromodomain inhibitor, bromosporine, which was previously co-crystallised with BD5.2, and the two compounds previously co-crystallised with BD5.1; SGC-CBP30 and BI 2536 (Figure S2; PDB codes 5TCM, 6BYA & 5TCK).

Thermal shifts were recorded for recombinant BD5.1, BD5.2 and BD5T with the 15 compounds (Figure S5). SGC-CBP30 binding increased the thermal stability of both BD5.1 (Δ*T*_m_ = 1.88 ± 0.24 °C when screened at 10 µM; Figure S6A) and BD5T (Δ*T*_m_ = 1.21 ± 0.06 °C when screened at 25 µM; Figure S6B), however, there was no indication of binding to BD5.2 (at 10 µM SGC-CBP30). Unexpectedly, bromosporine did not produce a positive increase in the melting temperature of BD5.1, BD5.2 or BD5T. Similarly, BI 2536 did not cause a positive thermal shift for BD5T and produced an insignificant increase in the thermal stability of BD5.1 (Δ*T*_m_ = 0.14 ± 0.23 °C when screened at 10 µM). This may indicate weak affinity binding of these compounds which does not stabilise the protein to such an extent that a significant thermal shift can be detected.

Of the remaining 11 compounds, most failed to significantly increase protein stability, however, one compound which emerged as another potential ligand of *Ld*BDF5 was the human BRD9 inhibitor, I-BRD9. This compound gave rise to thermal shifts for both BD5.2 (Δ*T*_m_ = 0.57 ± 0.05 °C when screened at 75 µM; Figure S6C) and BD5T (Δ*T*_m_ = 0.31 ± 0.07 °C when screened at 75 µM; 0.81 ± 0.05 °C when screened at 150 µM; Figure S6D). Whilst small, these thermal shifts were statistically significant (unpaired t-test, P < 0.01, n = 6), and as with the SGC-CBP30 thermal shifts, are clearly visible in the TSA melting curves (Figure S6). Thus, these results provide the first binding assay data for *Ld*BDF5 bromodomains interacting with bromodomain inhibitors.

### Investigating SGC-CBP30, Bromosporine and I-BRD9 as LdBDF5 Ligands

Based on both the TSA data and co-crystal structures, three compounds, SGC-CBP30, bromosporine and I-BRD9, were selected for further analysis. SGC-CBP30 (Figure 4A) was originally developed as an inhibitor of human CBP/p300, displaying nanomolar binding affinity (*K*_d_) to these bromodomains^54^. It contains a 3,5-dimethylisoxazole ring which acts as an acetyl-lysine bioisostere^53,55^. Bromosporine (Figure 4B), a broad-spectrum bromodomain inhibitor containing a triazolopyridazine dicyclic core, is a widely used tool compound which binds to a diverse range of bromodomains^56^. Finally, I-BRD9 (Figure 4C) is a thienopyridone derivative which binds to human BRD9, exhibiting 700-fold selectivity over the human BET bromodomains^57^.

**Figure 4.**
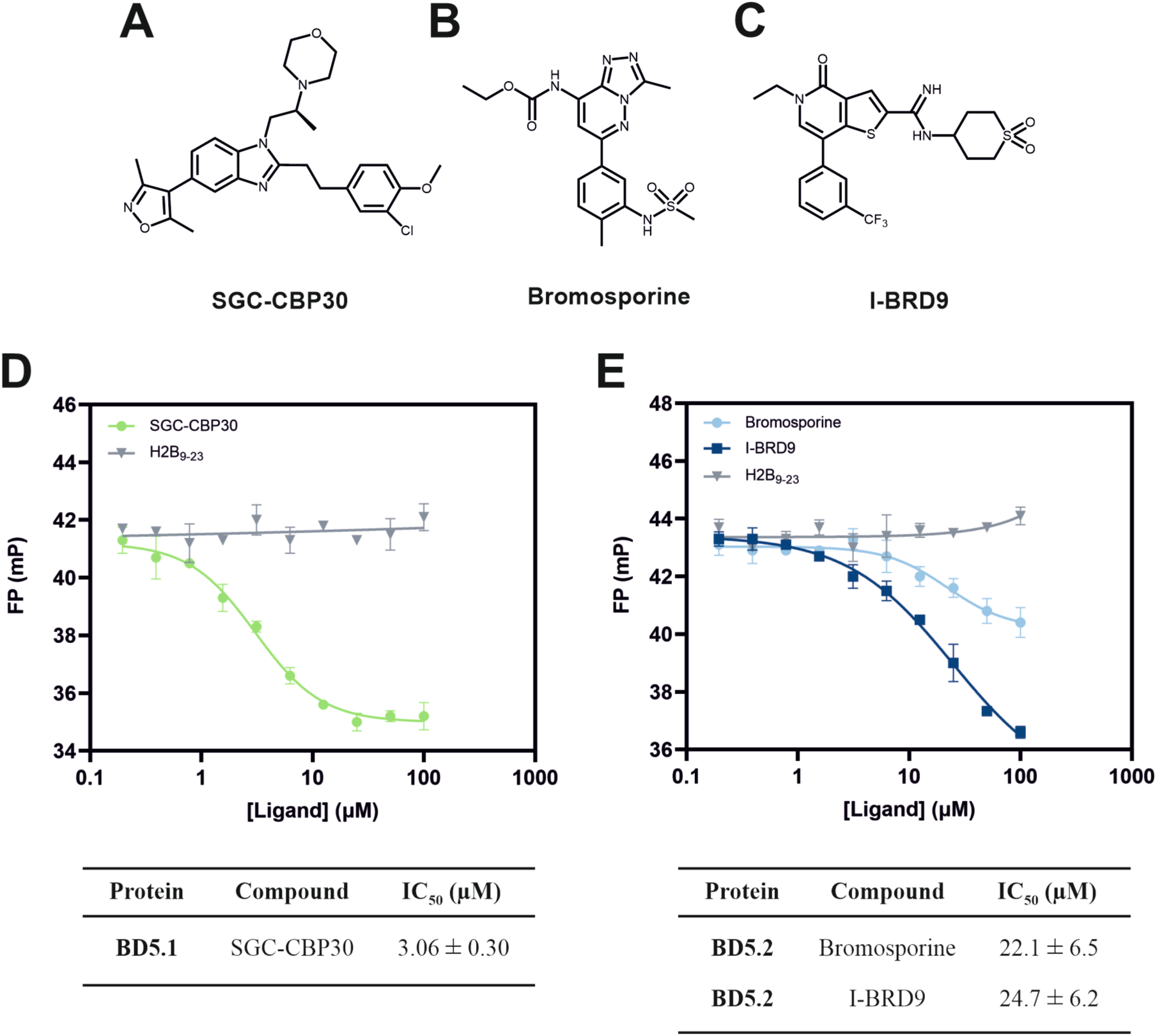
Structures of bromodomain inhibitors (A) SGC-CBP30, (B) bromosporine, and (C) I-BRD9. FP competition assays and associated IC_50_ values for displacement of the H2B_17-23_K19^Ac^K21^Ac^ probe by (D) SGC-CBP30 for binding to BD5.1, and (E) bromosporine and I-BRD9 for binding to BD5.2, compared with negative control unmodified unlabelled H2B_9-23_ peptide. In D and E mean, blank-corrected FP values are plotted against ligand concentration, and data fitted to [Inhibitor] vs. response -- Variable slope (four parameters) non-linear regression; error bars representing SD (n = 3).

First, competition binding assays were carried out to validate and quantify the binding of SGC-CBP30 to BD5.1, and bromosporine and I-BRD9 to BD5.2. Utilising the histone peptide that displayed the strongest binding to BD5.1 and BD5.2 (**H2B_17-23_K19^Ac^K21^Ac^**) as a probe, competitive binding FP experiments were performed. Compounds (0 – 100 µM) were titrated against fixed concentrations of the protein (2.5 µM BD5.1 or 10 µM BD5.2) and the fluorescent peptide probe (800 nM) (Figure 4D,E). When present at micromolar concentrations, all three compounds displaced the peptide probe, causing a reduction in FP. In comparison, an unmodified, unlabelled peptide based on the H2B_9-23_ sequence, included as a negative control, caused no probe displacement. The FP reduction elicited by bromosporine is anomalous, appearing to level off at a significantly higher mP than that of I-BRD9 which was closer to the expected FP for the free probe. Nevertheless, the BD5.2 ligands yielded similar IC_50_ values of 22.1 ± 6.5 and 24.7 ± 6.2 µM for bromosporine and I-BRD9, respectively. For the binding of SGC-CBP30 to BD5.1, an IC_50_ of 3.06 ± 0.30 µM was calculated. Though these values indicate low-affinity interactions, this was expected from non-optimised inhibitors and served to confirm the binding interactions. Additionally, the FP assay confirmed our prediction that the **H2B_17-23_K19^Ac^K21^Ac^** peptide binds in the bromodomain binding pocket, as it was displaced by compounds seen to bind within this binding cavity in co-crystal structures (Figure S2). A potential caveat to these assays is the low affinity (high *K*_d_) of the tracer binding the bromodomains, which can reduce the assay window and limit the identification of high-affinity inhibitors^58^. We, therefore, sought to increase our confidence in the binding of the compounds to the target bromodomains using orthogonal techniques, and potentially improve our estimate of their true potency.

Binding of the three compounds to *Ld*BDF5 bromodomains was next investigated using ligand-observed NMR. A combination of three experiments was used; waterLOGSY^59,60^, saturation transfer difference (STD)^61^, and CPMG^62,63^. Table 2 summarises the outcomes of each experiment. SGC-CBP30 showed evidence of binding to BD5.1 in all three experiments, with STD spectra informing on the binding epitope; peaks around 2.30 and 2.45 ppm, which can be assigned to the methyl groups of the 3,5-dimethylisoxazole group, showed markedly higher intensities than other peaks in the spectrum recorded for the compound with BD5.1 (Figure 5A). This correlates with the mode of binding of SGC-CBP30 observed in the co-crystal structure (Figure S2B), and with previous evidence that the 3,5-dimethylisoxazole group acts as an acetyl-lysine mimic, displacing acetylated histone peptides from bromodomains^53^. Binding of I-BRD9 to BD5.2 was supported by the results of CPMG and waterLOGSY experiments. In the waterLOGSY spectra, weak positive compound peaks were observed for I-BRD9 in the presence of protein, contrasting with the negative peaks in the compound alone spectrum, indicating compound binding (Figure 5B). The STD experiment for I-BRD9 was less conclusive, with most compound peaks either absent or very weak, with the exception of those at higher chemical shifts around 8.0 – 8.3 ppm, however, overall, these results were concordant with binding to BD5.2. For bromosporine, there was no evidence of binding to BD5.2 from the STD or CPMG spectra, while the waterLOGSY experiment produced ambiguous results.

**Figure 5.**
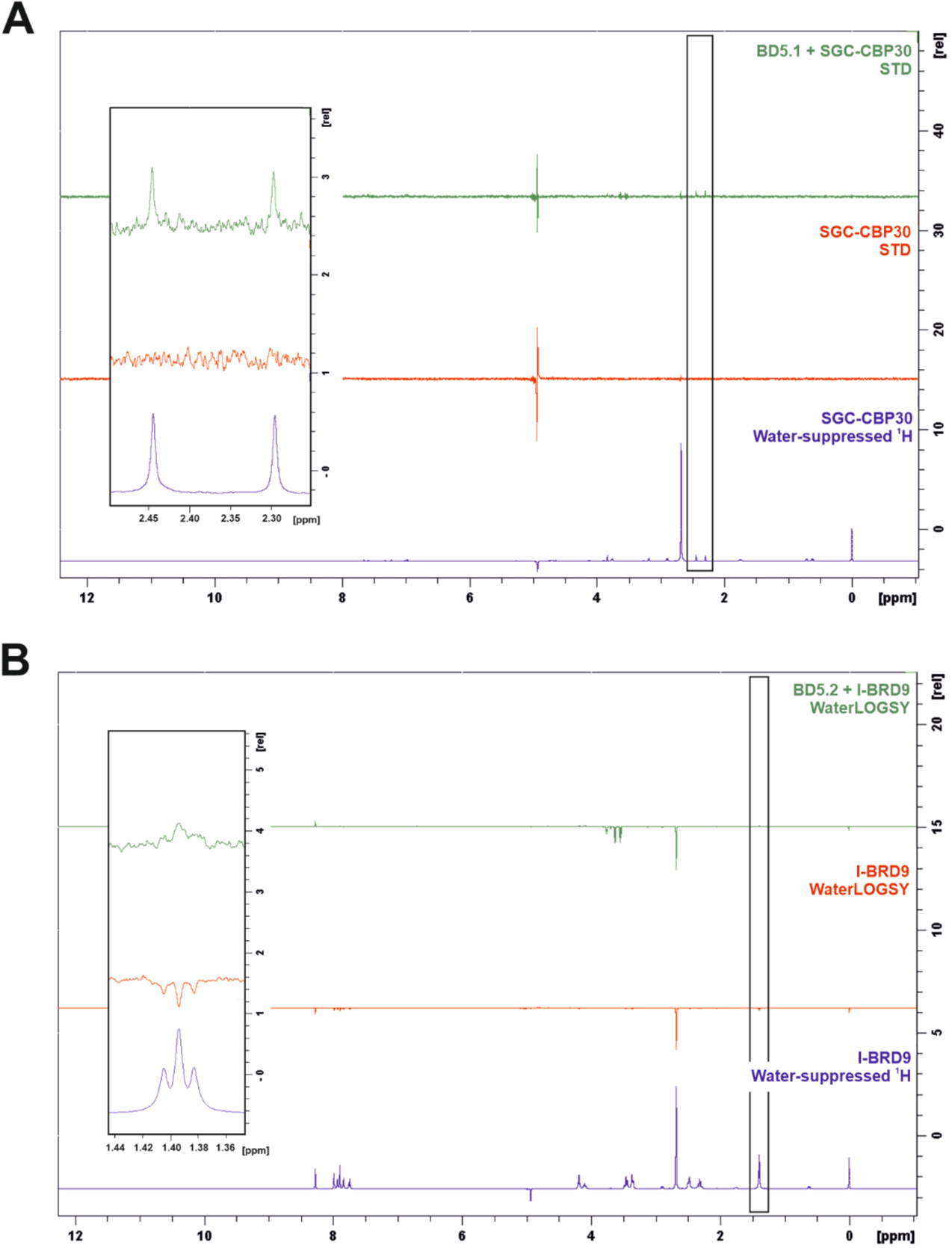
(A) STD NMR spectra recorded for SGC-CBP30 in the absence (red), and presence (green) of BD5.1, alongside the reference water-suppressed ^1^H spectrum for the compound alone (blue); positive peaks in the STD spectra indicate ligand binding to the protein. (B) WaterLOGSY NMR spectra recorded for I-BRD9 in the absence (red), and presence (green) of BD5.2, alongside the reference water-suppressed ^1^H spectrum for the compound alone (blue); representative compound peaks are indicated on the full spectra and shown as an inset.

**Table 2.** Outcomes of NMR experiments for BD5.1 and BD5.2 binding bromodomain inhibitors, where tick marks represent positive confirmation of binding, question marks indicate inconclusive results and crosses indicate results which did not suggest a binding interaction.

| Protein | Compound | WaterLOGSY | STD | CPMG |
| --- | --- | --- | --- | --- |
| BD5.1 | SGC-CBP30 | ✓ | ✓ | ✓ |
| BD5.2 | Bromosporine | ? | × | × |
| BD5.2 | I-BRD9 | ✓ | ? | ✓ |

We next quantified the BD5.1 affinity of SGC-CBP30 using microscale thermophoresis (MST) and isothermal titration calorimetry (ITC) (Figure 6). SGC-CPB30 was determined to have a *K*_d_ value for BD5.1 of 281 ± 12.3 nM in ITC. We were able to corroborate this result in the MST assay which determined binding of the compound with a *K*_d_ value of 369 ± 31.6 nM. SGC-CBP30 has high affinity for BD5.1 and could therefore provide a good starting point for developing ligands and tools to study BD5.1. As mentioned, the FP assay is limited by the relatively low-affinity binding of the fluorescent peptide probe, therefore the IC_50_ values calculated using this method are likely higher than their true values^58^. Use of ITC and MST has enabled us to more accurately determine *K*_d_ values for SGC-CBP30 for BD5.1, demonstrating that this compound is a high-affinity ligand for the protein module.

**Figure 6.**
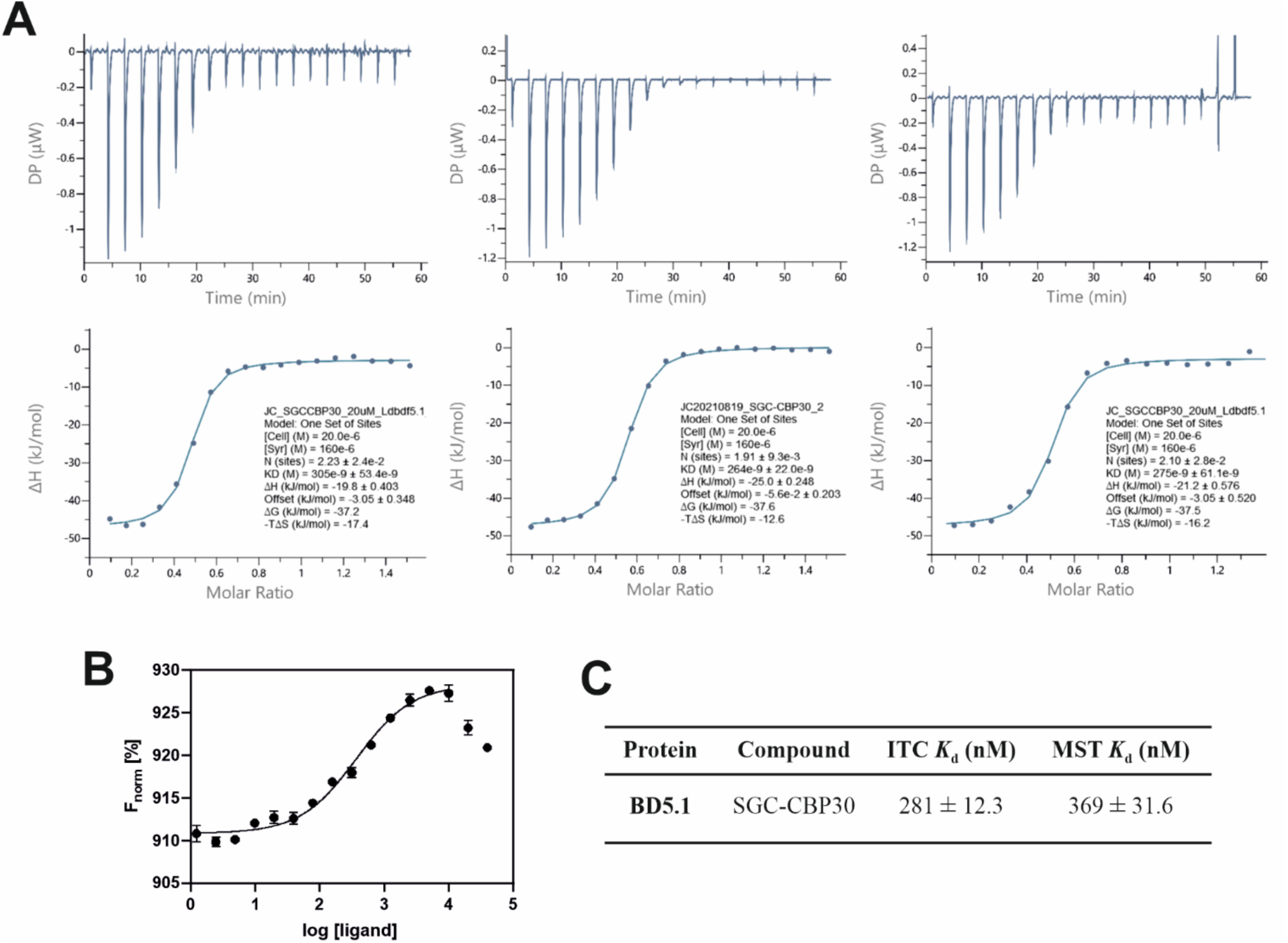
(A) Isothermal titration calorimetry (ITC) data for SGC-CPB30 (20 μM, in the cell), measured with BD5.1 (160 μM, in the syringe). Shown are heat effects for each injection (above) and the normalised binding isotherms (below) including the fitted function for the compound binding (solid line). (B) Microscale thermophoresis binding curve for SGC-CPB30 in BD5.1, calculated from the gradual difference of thermophoresis between the fluorescent molecules of both unbound and bound states, which is plotted as F_norm_ (defined as F_hot_/ F_cold_) against ligand concentration. Graph produced in GraphPad Prism with data fitted to log(agonist) *vs*. response; error bars representing SEM (n = 3). (C) Associated *K*_d_ values for ITC and MST; for ITC *K*_d_ was determined from analysis in MicroCal PEAQ-ITC Analysis software (Malvern 1.1.0.1262) using a single binding site model; for MST *K*_d_ was derived from the binding curve.

### Quantification of Bromosporine Solubility

The binding assay data for bromosporine with BD5.2 described above produced ambiguous results, with TSA and NMR failing to detect a clear binding interaction, and the FP competition curve plateauing at an unexpected FP value. These results were surprising as a crystal structure of BD5.2 in complex with bromosporine has been solved (PDB code 5TCK). We considered therefore whether our experiments might have been compromised by incomplete dissolution of bromosporine. Bromosporine solubility was quantified using a Chromatographic logD (ChromlogD) assay^64^ which measures the lipophilicity of a compound. In this chromatography-based assay, the column retention time of a compound of interest is measured and correlated with a chromatographic hydrophobicity index (CHI) and subsequently projected to a logP/D scale. The chromlogD_pH7.4_ value for bromosporine was calculated to be 3.34 which indicates moderate to low solubility. SGC-CBP30 and I-BRD9 were calculated to have a chromlogD_pH7.4_ value of 5.69 and 3.93 respectively.

### Probing the Interaction of Bromosporine with BD5.2 using X-ray Crystal Structures

Further analysis revealed an anomalous mode of ligand binding in the BD5.2-bromosporine co-crystal structure, at least in the crystal form. Crystals of bromosporine-bound (Figure 7A) and unliganded (Figure 7B) BD5.2 are isomorphous and the two chains in the asymmetric units can be overlaid using SSM superpose routines to give an rmsΔ value of 0.65 Å for 272 equivalent atoms. This close similarity of packing was unexpected because the bromosporine ligands appeared to be important determinants of the molecular packing observed in the co-crystal structure. As can be seen in Figure 7A, the pair of bromosporine molecules contribute significantly to the interface between the A and the B molecules of the asymmetric unit. Analysis of the molecular interfaces in the program PISA^65^, shows that of the 630 Å^2^ of surface area on each ligand, 200 Å^2^ is buried in the interface with chain A, 195 Å^2^ is buried in the interface with chain B and 130 Å^2^ is buried in bromosporine-bromosporine interactions. Thus, each of the bromosporine ligands is effectively shared by the A and B chains.

**Figure 7.**
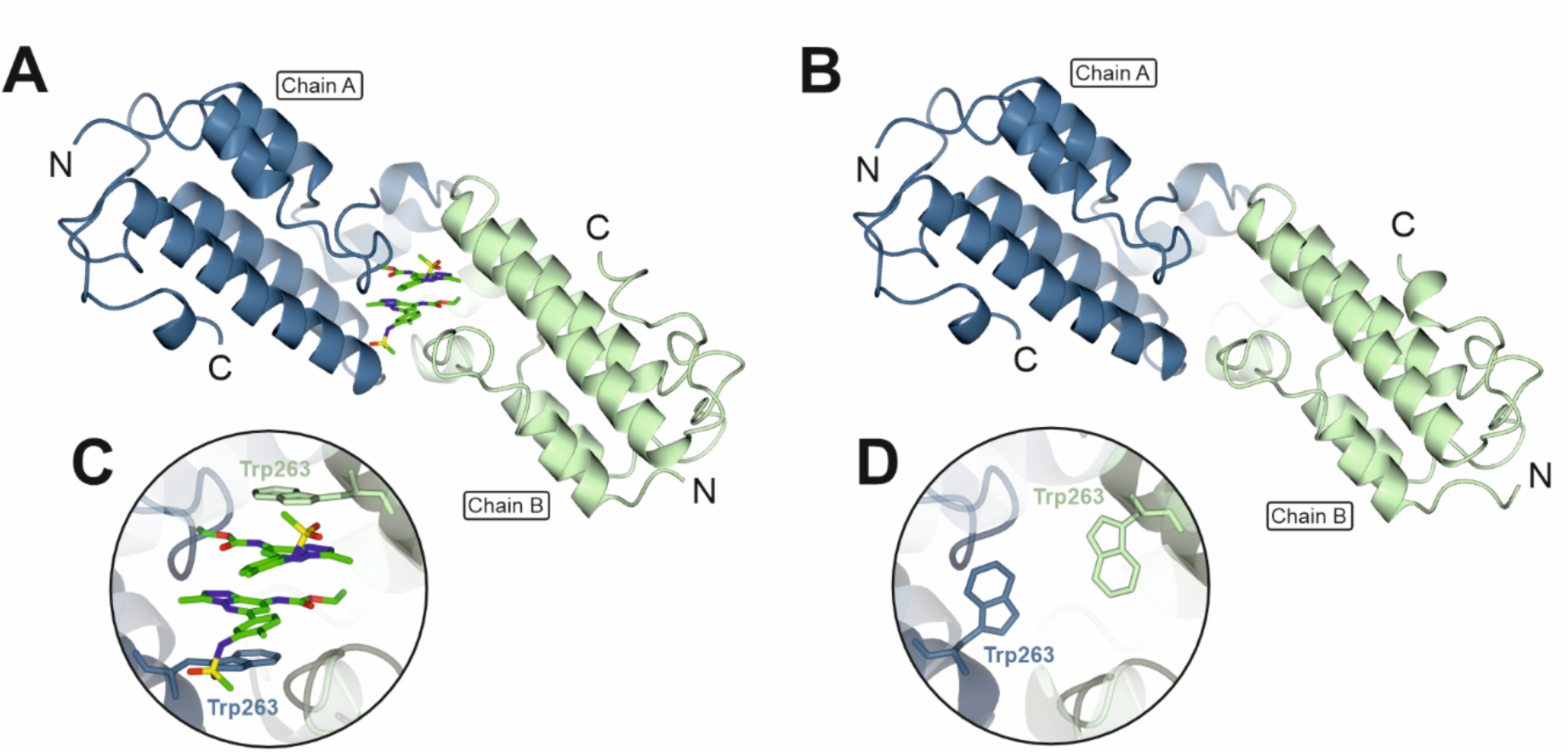
X-ray crystal structures of BD5.2, showing A and B chains of the (A) bromosporine-bound (PDB code 5TCK) and (B) unliganded (PDB code 8BPT) bromodomain asymmetric units, with (C, D) insets showing movement of Trp263 accompanying bromosporine binding.

The clearest difference between the liganded and unliganded BD5.2 structures results from the implied movement of W263 accompanying the binding of bromosporine (Figure 7C,D). In the complex, the pair of ligands form aromatic ring stacking interactions with one another and with the flanking indole rings of W263. In the absence of bromosporine, the side chains of W263 adopt different rotameric conformations and partially occupy the volume taken up by the compound. Significant conformational changes of the surrounding aromatic side chains of Y201 and F256 also accompany bromosporine binding.

In crystal structures of various human bromodomains, as well as the bromodomains of *Ld*BDF2 and *Ld*BDF3, bromosporine binds in the acetyl-lysine binding pocket with the core bicyclic ring oriented such that its methyl substituent points furthest into the deep cavity (Figure S7). The nitrogen of the pendant ethyl carbamate substituent, together with a nitrogen from the triazole ring, form hydrogen bonds to the side chain amide of the highly conserved Asn from the BC loop. Meanwhile, the sulfonamide-containing side chain projects along a groove formed by the ZA loop. The mode of bromosporine binding in the BD5.2 complex is very different, with the plane of the triazolopyridazine moiety almost perpendicular to its orientation in the other bromodomains, such that the sulfonamide and ethyl carbamate moieties project in different directions (Figure S7). There is very little overlap in the volume occupied by the bromosporine ligand in the *Ld*BDF5 BD5.2 crystal structure and the volumes occupied by the ligands in the structures of the other bromodomain-bromosporine complexes. The conformation of Trp263 in the unliganded BD5.2 structure is similar to that of the corresponding Trp93 in the BD2-bromosporine crystal structure. In the other orthologous complexes presented in Figure S7, Trp263 is replaced by Tyr and Ile residues, and the side chains of these residues also pack against the face the bicyclic ring in bromosporine. Overall, we suggest that the bromosporine binding to BD5.2 in the crystal structure is an artefact of crystal packing and that bromosporine is, at best, a low-affinity ligand of this domain.

### Activity of Human Bromodomain Inhibitors against Leishmania Promastigotes

Whilst genetic validation studies have established that BDF5 is essential in *Leishmania*^14^, and biophysical assays here demonstrated its ability to be targeted by bromodomain inhibitors, it was also important to ascertain whether bromodomain inhibitors inhibit the growth of the parasite. To this end, we performed cell viability assays with SGC-CBP30, bromosporine and I-BRD9. These compounds were screened against *L. mexicana* and *L. donovani* promastigotes as genetic target validation of BDF5 in *L. mexicana* indicated it is an essential protein and the bromodomains are highly conserved in the different species^14^. Therefore, effective bromodomain inhibitors have the potential to have broad anti-leishmania activity. Parasites were exposed to the compounds (0 – 30 µM) in technical triplicates for five days, before measuring cell viability using resazurin. Mean fluorescence intensity measurements were normalised to give values as percentage cell viability, averaged over triplicate biological replicates, and EC_50_ values calculated. Bromosporine and I-BRD9 showed minimal activity up to 30 µM, however, SGC-CBP30 exhibited moderate cytotoxicity towards both species, with EC_50_ values approaching that of the established antileishmanial, miltefosine (Figure 8). Despite SGC-CBP30 not being optimised against *Leishmania* bromodomains, its activity indicates a promising starting point for exploring medicinal chemical optimisation of more potent binders of BD5.1. Genetic approaches indicated a level of redundancy between BD5.1 and BD5.2 in *Ld*BDF5^14^ but there is precedent of potent binders of single bromodomain in multi-bromodomain proteins having phenotypic effects ^66^. A highly potent binder could also be used as a starting point for a PROTAC strategy to degrade BDF5 and its associated proteins^67^, however, this is caveated by the current lack of validated ubiquitin ligase binding compounds that can recruit parasite E3 ligases to target proteins. Other hydrophobic tagging groups could also be investigated as destabilising agents to trigger selective BDF5 degradation^68^.

**Figure 8.**
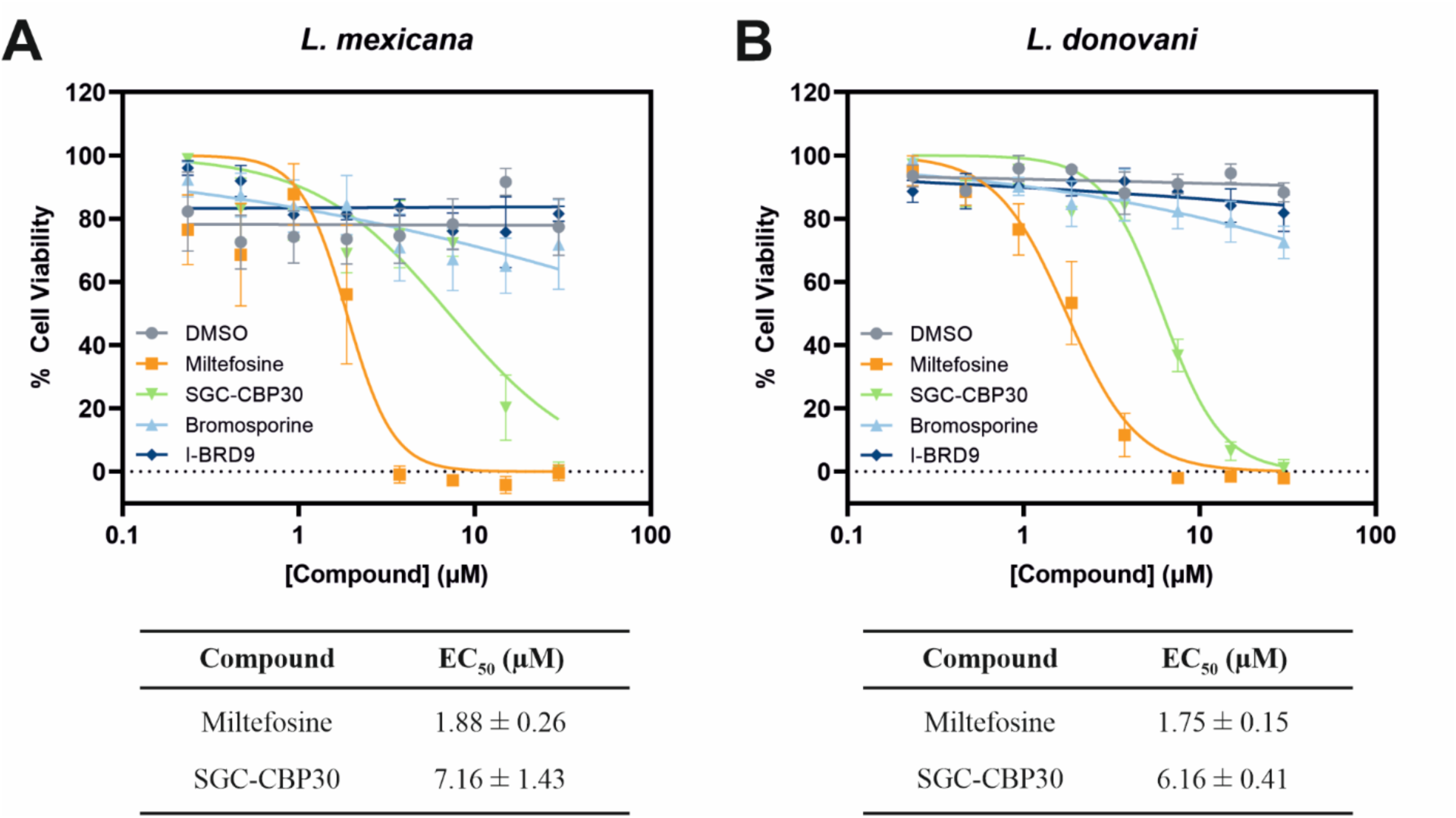
Dose-response curves for cell viability assays after five-day incubation of human bromodomain inhibitors and miltefosine with (A) *L. mexicana* and (B) *L. donovani* promastigotes. Mean, blank-corrected fluorescence intensity measurements were normalised and averaged over three biological replicates; fitted to an [Inhibitor] *vs*. normalized response -- variable slope dose-response model for EC_50_ calculations. Error bars represent SEM (n = 3).

## Conclusions

Regulation of gene expression in *Leishmania* is complex and tightly controlled, allowing the parasite to differentiate and adapt to different environments at appropriate points in its lifecycle. Research is now beginning to shed light on how epigenetics maps onto this landscape, with indications that reader proteins such as BDF5 can regulate global RNA polymerase II transcription^14^. Whilst the precise mechanism by which BDF5 exerts this effect remains to be elucidated, we have identified interactions with peptides from two histone proteins, H2B and H4, where the latter binding sequence contains known HAT acetylation sites^44–48^. Both histone peptides bound to both BD5.1 and BD5.2, hinting at a cooperative mechanism of binding. Whilst weak, the observed binding affinities are comparable with those reported for human bromodomains binding mono-acetylated histone peptides measured in a large-scale analysis^19,69,70^. Furthermore, BDF5 has the potential to bind chromatin as part of the CRKT complex^14^. This multiprotein complex is also predicted to contain bromodomain proteins BDF3 and BDF8, thus multivalency may increase the bromodomain binding avidity and specificity.

Although the biological relevance of the acetylated H2B sequence is not yet established, the peptides described in this work represent tool compounds, applied here in a bespoke FP assay. Applying this technique alongside orthogonal assays we examined the potential BDF5 holds as a target for antileishmanial drug discovery. We report three human bromodomain inhibitor compounds which bind to the BDF5 bromodomains in orthogonal biophysical assays; SGC-CBP30, bromosporine and I-BRD9. Encouragingly, the compound SGC-CBP30, which binds to BD5.1 (*K*_d_ = 281 ± 12.3 nM as measured by ITC) also exhibited growth inhibition towards *Leishmania* promastigotes in cell viability assays, consistent with previous evidence that this protein is essential for the parasite^24^. SGC-CBP30 contains the known 3,5-dimethylisoxazole acetyl-lysine mimic, which has been shown to prevent histone binding to human bromodomains. STD NMR data indicated that this moiety contributes to the binding epitope, consistent with a canonical mode of bromodomain inhibition^53,55^. With the proviso that the *in vivo* properties of SGC-CBP30, such as potential off-target binding, need to be further investigated, this compound provides a useful starting point for the development of more potent inhibitors of BDF5. Moreover, this work highlights the chemical tractability of *Leishmania* BDF5 and provides a strong rationale for future exploration of BDF5 inhibitors in antileishmanial drug discovery.

## Methods

### Recombinant Protein Expression and Purification

Based upon native protein sequences of *Ld*BDF5 (LDBPK_091320) and *Ld*BDF2 (LDBPK_363130), plasmids containing the bromodomain cassettes *Ld*BDF5 BD5.1, BD5.2, BD5T and BDF2 (Table S1–4) were used for recombinant expression of these proteins. Plasmids were kindly gifted by Dr Raymond Hui at the Structural Genomics Consortium, Toronto, using vectors derived from a pET-15-MHL backbone plasmid, produced by Dr Cheryl Arrowsmith at the Structural Genomics Consortium (Addgene plasmid #26092). The pET-15 plasmid allows for IPTG-induced protein expression under the control of a T7 promoter. The plasmid also carries an ampicillin resistance gene and encodes a cleavable His-tag.

For the expression of *Ld*BDF2 and *Ld*BDF5 BD5T bromodomains, overnight cultures of *E. coli* RosettaTM (DE3) pLysS cells (carrying chloramphenicol resistance) harbouring the relevant plasmids were used to inoculate 0.5 – 1 L Luria-Bertani broth (LB) supplemented with ampicillin (100 µg/ml), chloramphenicol (35 µg/ml) and glucose (0.2%) and grown at 37 °C with shaking at 180 rpm. Once the OD_600_ reached 0.6, recombinant protein production was induced by adding IPTG (to 1 mM), followed by incubation at 37 °C for 2.5 hours for *Ld*BDF2, or 30 °C for 3.5 hours or overnight for *Ld*BDF5 BD5T. Cells were harvested by centrifugation at 5,000 rpm 4 °C for 30 mins. Cell pellets were resuspended in a lysis buffer (20 mM HEPES, 500 mM NaCl, 30 mM imidazole, pH 7.5) at 5 ml per 1 g cell pellet, and the suspension supplemented with 1 cOmplete^TM^ mini EDTA-free protease inhibitor cocktail tablet (Roche), MgCl_2_ (5 mM) and DNase I (5 μg/ml).

Lysis was performed by sonication on ice using 30 second bursts separated by 30 second pauses for a total of 15 minutes and cell debris pelleted by centrifugation at 15,000 rpm, 4 °C for 40 minutes. The soluble lysate was resolved by IMAC with an ÄKTA pure protein purification system (Cytivia). The sample was applied to a 5 ml HisTrap^TM^ FF column (Cytivia), which was developed with a linear gradient of low (20 mM HEPES, 500 mM NaCl, 30 mM imidazole, 1 mM DTT pH 7.5) to high (20 mM HEPES, 500 mM NaCl, 300 mM, imidazole, 1 mM DTT, pH 7.5) imidazole buffer over 10 column volumes (CV) at a flow rate of 2 ml/min. Fractions were analysed by SDS-PAGE and those containing the protein of interest were combined and concentrated. Cleavage of the His-tags was achieved by addition of recombinant tobacco etch virus (TEV) protease at 1 mg per 50 mg protein, alongside dialysis into imidazole-free buffer (20 mM HEPES, 500 mM NaCl, pH 7.5). The proteins were then concentrated and applied to a second HisTrap^TM^ column, washed with 5 CV low imidazole buffer and subsequently developed with a linear gradient of low to high imidazole buffer over 20 CV at 2 ml/min. Fractions were again analysed by SDS-PAGE, pooled and concentrated. Further purification was achieved using a HiLoad 16/600 Superdex 75 pg size exclusion chromatography column with an ÄKTA pure protein purification system (both Cytivia). Protein samples (< 2.5 ml) were loaded onto the column and the column washed with 1 CV (120 ml) imidazole-free buffer. Fractions were pooled based upon SDS-PAGE analysis and concentrated.

Production of recombinant *Ld*BDF5 BD5.1 and BD5.2 proteins was performed by batch-culture using 7 L glass bioreactors (Applikon). Plasmids were transformed into Rosetta^TM^2 (DE3) pLysS competent cells and used to inoculate overnight starter cultures of LB and ampicillin (100 µg/ml), incubated at 37 °C, 200 rpm for 16 hours. 50 ml culture was then used to inoculate bioreactors containing 5 L LB and ampicillin (100 µg/ml), with bioreactors configured for cultivation at 37 °C, 2 L/min air-flow, dO_2_ maintained at 30% (saturation in air) using a stirrer cascade. Once the OD_600_ reached 1.0, temperature was lowered to 18 °C for 30 minutes after which time IPTG was added (0.1 mM). Cultures were further incubated for 20 hours and cells harvested by centrifugation at 7,000 rpm 4 °C for 10 mins. Cell pellets were resuspended in a lysis buffer (20 mM HEPES, 500 mM NaCl, 10 mM imidazole, 5% glycerol) at 5 ml per 1 g cell pellet, supplemented with DNAse I (4 U/ul) and 1 cOmpleteTM mini EDTA-free protease inhibitor cocktail tablet (Roche). The sample was lysed by French press at 17 KPsi (Constant Systems Ltd), and any remaining sample washed with 15 ml lysis buffer, with the lysate centrifuged at 17,000 x g, 4°C for 10 minutes. The samples were syringe filtered and loaded onto a 5 ml HisTrap^TM^ FF crude column (Cytivia) with a BioRad NGC Chromatography system, developed using a linear gradient of low imidazole buffer (20 mM HEPES, 500 mM NaCl, 30 mM imidazole, 5% glycerol) to high imidazole buffer (20 mM HEPES, 500 mM NaCl, 500 mM imidazole, 5% glycerol) over 10 CV. Fractions were analysed by SDS-PAGE, and those containing protein pooled and concentrated. His-tags were not cleaved for BD5.1 and BD5.2 recombinant proteins. Further purification was achieved by loading concentrated 2 ml samples onto a HiLoad 16/600 Superdex 75 pg size exclusion chromatography column (Cytivia) with a NGC chromatography system (BioRad), eluted using an imidazole-free buffer (20 mM HEPES, 500 mM NaCl, 5% glycerol). Fractions were analysed by SDS-PAGE, pooled and concentrated as before. All protein concentrations were calculated by measuring absorbance at 280 nm (A_280_), with relevant the molar extinction coefficient determined by the Protparam online tool within the Expasy Swiss bioinformatics resource portal.

### SEC-MALLS

Purified recombinant proteins in HEPES buffer (20 mM HEPES, 500 mM NaCl, pH 7.5) were injected in 100 μl volumes onto a Superdex S75 size exclusion column (Cytivia), run at 0.5 ml/min using an HPLC system (Shimadzu). Data were collected using an HPLC system SPD-20A UV detector (Shimadzu) and a HELEOS-II multi-angle light scattering detector and rEX refractive index detector (both Wyatt). Data analysis was performed using Astra 7 software and protein molecular mass estimated by the Zimm fit method with degree 1. A control sample of BSA was run to correct for changes in instrument calibration and dn/dc values in the buffer used. Graph plotted using GraphPad Prism® software.

### X-ray Crystallography

Crystals of BD5.2 were obtained from solutions containing 0.1 M HEPES pH 7.5 and 1.4 M sodium citrate. Data extending to 1.6 Å were collected from a single crystal on beamline i04 at the DIAMOND Light Source. The data were processed in xia2-3d^71^ revealing that the crystals belong to space group P2_1_2_1_2_1_ with cell dimensions of 33.4 Å, 75.4 Å and 105.8 Å indicating the presence of two chains in the asymmetric unit of the crystal and a solvent content of 42%. Data collection and refinement statistics are given in Supplementary Table S5. The structure was solved in MOLREP^72^ using chain A of the coordinates of the BD5.2– bromosporine complex (PDB code 5TCK) as the search model. Two clear solutions were obtained consistent with expectations, and giving R_work_/R_free_ values of 0.33 and 0.36, respectively. The structure was refined using cycles of REFMAC5^73^ interspersed with manual modelling in COOT^74^ with the introduction of 147 water molecules. TLS and anisotropic temperature factor refinement were included in the later refinement cycles yielding final R_work_ and R_free_ values of 0.17 and 0.23, respectively. The fit to the electron density maps is good with the exception of Arg180 in both chains and Glu300 in chain A.

### Histone Peptide Microarray

*L. donovani* histone sequences were identified using TriTrypDB (PMID: 36656904) and the first 25 amino acids after the start methionine were selected for all non-redundant sequences. Histone H2AZ was omitted from the array due to the extended size of the N terminal tail. These 25mer sequences were submitted to PepperPrint (Heidelberg) for tiling onto a glass slide using a solid-phase fmoc chemistry laser printer. Peptides were 15-mers that were shifted by 1 amino acid per spot and tiled to span the 25-mer. A section of the array contained the unmodified amino acid sequence, a section contained peptides where all lysine residues were replaced with acetyl-lysine, and another section contained peptides where only single lysine residues were replaced by acetyl-lysine. Peptides were spotted in duplicate. The array was probed with a modified version of the PepperCHIP immunoassay protocol. The array was wetted with PBST (PBS pH 7.4, 0.05% Tween 20) and then blocked with PBST 1% BSA for 30 min at room temperature. Recombinant proteins were diluted to 5 mg/ml in PBST without BSA overnight at 4 °C with rotational mixing at 140 rpm. The array was then washed twice with PBST for 10 seconds. Conjugated primary antibodies (monoclonal mouse anti-HA (12CA5) Cy5 control and 6X His Tag Antibody Dylight™ 549 Conjugated) were applied to the array at 1:500 dilution in PBST 0.1% BSA for 30 mins at room temperature. The array was washed twice with PBST for 10 seconds each time, dipped 3x 1 sec in 1 mM Tris pH 7.4, dried with a stream of compressed air and then imaged using an Agilent DNA microarray scanner. The fluorescent intensity of each spot was quantified using PepSlide Analyzer software. Graph plotted using using GraphPad Prism® software.

### Histone Peptide Design and Synthesis

Peptides were designed based on *L. donovani* histone H2B (LDBPK_171320) and histone H4 (LDBPK_150010) sequences. Peptide variants were designed including acetylated versions and unmodified control peptides, with a conjugated 5-carboxyfluorescein (5-FAM) fluorophore (excitation and emission of around 492 and 518 nm, respectively). The fluorophore was covalently joined either to the N-terminus or via an ethylene diamine linker to the C-terminus. Unlabelled versions were also produced for use in TSA or as control peptides. Peptides were synthesised by Cambridge Research Biochemicals at > 90% purity with analysis by HPLC and mass determination by MALDI coupled to time-of-flight (MALDI-TOF) mass spectrometry. Peptides were supplied as 5 mg or 10 mg quantities (Cambridge Research Biochemicals) and dissolved to 10, 50 or 100 mM in dimethyl sulfoxide (DMSO).

### Human Bromodomain Inhibitor Compounds

Bromosporine, I-BET151, SGC-CBP30, BI 2536 and JQ1 were purchased commercially from Advanced ChemBlocks, Cayman Chemical and Sigma-Aldrich. GSK8814 was supplied by the Structural Genomics Consortium under an Open Science Trust Agreement. Compound stock solutions were prepared to concentrations of 10 – 100 mM in DMSO, or deuterated DMSO to allow them to also be used in the NMR assay.

### Thermal Shift Assay (TSA)

Following TSA optimisation experiments, concentrations were established for BD5.1 (3 µM protein; 3x dye), BD5.2 (3 µM protein; 2x dye), and BD5T (2.1 µM protein; 2x dye), using SYPRO orange dye (Merck). Dilutions were carried out in HEPES buffer (20 mM HEPES, 500 mM NaCl, pH 7.5). Control experiments were performed to confirm that compounds and peptides did not give fluorescent signals themselves or interfere with the assay. 25 μl samples were prepared containing the protein and dye plus ligand or DMSO for reference samples. Samples were dispensed into 96-well qPCR plates (Agilent) in replicates of six. Triplicate control samples were also included of the protein alone and dye alone. Plates were sealed and centrifuged at 2,000 rpm, 4 °C for 1 minute, and TSA experiments carried out using a Stratagene Mx3005P real-time PCR instrument (Agilent), increasing the temperature from 25 – 95 °C at 30 seconds per degree, taking fluorescence readings after each 30 second increment. Data were imported into an online JTSA tool for analysis (http://paulsbond.co.uk/jtsa), with a five-parameter sigmoid equation curve fitting applied to the data and melting points calculated as the midpoints^75^. Anomalous results within the six replicates were excluded from analysis where atypical melting curves were observed or low R^2^ values were generated for curve fitting. Melting temperatures were analysed in Microsoft Excel and GraphPad Prism®, with thermal shifts calculated as the difference between the melting temperature (*T*_m_) of the protein-ligand samples and the reference samples, reported as Δ*T*_m_ ± standard deviation. Where appropriate, statistical analysis was performed using an unpaired t-test.

### Fluorescence Polarisation (FP)

FP experiments were set up in 384-well black flat bottom plates (Corning) using sample volumes of 20 μl, diluting in FP buffer (20 mM HEPES, 500 mM NaCl, 1 mg/ml BSA added fresh, pH 7.5). Fluorescence intensity and polarisation readings were taken using a BMG LABTECH CLARIOstar® microplate reader following gain and focus adjustment. Data were processed using BMG LABTECH Mars software with subsequent analysis and graphs plotted using Microsoft Excel and GraphPad Prism®. Probe optimisation was performed using triplicate samples of 0 – 1000 nM probes alongside triplicate control samples of buffer alone for blank correction. Fluorescence intensity and polarisation readings were blank-corrected and FP calculated using the equation:

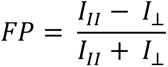

Here, I_II_ is intensity of emitted light polarised parallel to the excitation light, and I_⊥_ is intensity of emitted light polarised perpendicular to the excitation light. Mean, blank-corrected fluorescence intensity and polarisation values were plotted against probe concentration (Figure S8). 800 nM was deemed an appropriate concentration of probes to use in subsequent experiments. Protein binding experiments were then performed with samples containing the probe (800 nM) and protein (covering 0 – 300 µM) alongside control samples of probe alone and buffer alone, all samples prepared in triplicate. Samples were mixed in the wells, and the plate incubated in the dark for 30 minutes at room temperature before taking FP readings. Mean, blank corrected FP values were transformed by subtracting mP of the free probe, with ΔmP plotted against protein concentration and for BD5.1 and BD5.2, fitted to a one site-specific binding non-linear regression model for *K*_d_ determination. For the BD5T, a two sites-specific binding non-linear regression model was instead used.

The **H2B_17-23_K19^Ac^K21^Ac^** peptide was used as a probe in competition assays at 800 nM with 2.5 µM BD5.1 or 10 µM BD5.2. Fixed concentrations of the protein and probe were titrated against increasing concentrations of the competitor ligand up to 100 µM, obtained by serial dilution in DMSO (final concentration 1% DMSO). Protein and compounds were incubated together for 30 minutes at room temperature before addition of the probe, then FP readings taken as before. Controls included buffer alone, probe alone, protein alone and ligands alone. Mean, blank-corrected FP values were plotted against compound concentration and IC_50_ values calculated by fitting an [Inhibitor] vs. response -- Variable slope (four parameters) non-linear regression model.

### Ligand-observed Nuclear Magnetic Resonance (NMR)

Three different ligand-observed proton NMR experiments were performed; water ligand-observed via gradient spectroscopy (waterLOGSY), saturation transfer difference (STD), and Carr-Purcell-Meiboom-Gill (CPMG). Sodium Trimethylsilylpropanesulfonate (DSS) was used as the NMR reference standard to provide the standard peak, set to chemical shift 0 ppm. Proteins were transferred into a sodium phosphate (NaPi) NMR buffer (20 mM NaPi, 100 mM NaCl, pH 7.5) by buffer exchange and bromodomain inhibitor compound stocks were prepared in deuterated DMSO. Samples (550 μl) were prepared in 5 mm NMR tubes (Wilmad) including the appropriate controls, containing the protein (20 μM) and compound (0.6 mM) alongside D_2_O (13 or 17.5%), DSS (80 μM), sodium phosphate (20 mM) and NaCl (100 mM). Spectra were recorded using a 700 MHz Bruker Avance Neo spectrometer, equipped with a cryoprobe at 298 K, with 16 – 24 scans. 1D proton spectra were recorded with water suppression, alongside the three ligand-observed experiments. Spectra were analysed using TopSpin NMR data analysis software (Bruker) and proton chemical shifts predicted using the NMRDB NMR spectral predictor tool^76,77^. In waterLOGSY, a positive result was characterised by opposite signal peaks for the compound compared with non-binding compounds. In STD, saturation is transferred from the protein to bound compounds, therefore a positive result was recorded where compound peaks were observed in the ‘difference’ spectra. In CPMG, binding compounds exhibit a reduction in peak intensity with a longer relaxation delay, so when comparing spectra for compound alone with compound + protein, a greater reduction in peak intensity in the presence of the protein indicates a positive result.

### Isothermal Titration Calorimetry (ITC)

All calorimetric experiments were performed on a MicroCal PEAQ-ITC Automated (Malvern) and analysed with the MicroCal PEAQ-ITC Analysis software (Malvern 1.1.0.1262) using a single binding site model. The first data point was excluded from the analysis. The BD5.1 bromodomain was dialysed at 4 °C overnight in a Slide-A-Lyzer® MINI Dialysis Device (2000 MWCO; Thermo Scientific Life Technologies) into HEPES buffer (20 mM HEPES, 500 mM NaCl, pH 7.4) containing 0.5% DMSO. Proteins were centrifuged to remove aggregates (2 min, 3,000 rpm, 4 °C). Protein concentration was determined by measuring the absorbance at 280 nm using a NanoDrop Lite spectrophotometer (Nanodrop® Technologies Inc.) by using the predicted protein absorbance (*Ld*BDF5.1: ε280: 12800 M^−1^ cm^−1^). The ligand was dissolved as 20 mM DMSO stock solution and diluted to the required concentration using dialysis buffer. The cell was stirred at 750 rpm, reference power set to 5 μcal/sec and temperature held at 25 °C. After an initial delay of 60 sec, 20 × 2 μl injections (first injection 0.4 μl) were performed with a spacing of 150 sec. Heated dilutions were measured under the same conditions and subtracted for analysis. Small molecule solutions in the calorimetric cell (250 μl, 20 μM) were titrated with the protein solutions in the syringe (60 μl, 160 μM).

### Microscale Thermophoresis (MST)

The BD5.1 protein was labelled with the *Monolith His*-*Tag Labeling Kit* (RED-tris-NTA 2nd Generation *kit*, NanoTemper Technologies, Catalog# MO-L018) according to the manufacturer’s protocol. The test compound was serially diluted in 16-steps in PBS-T buffer (137 mM NaCl, 2.7 mM KCl, 10 mM Na_2_HPO_4_, 1.8 mM KH_2_PO_4_, 0.1% Tween-20, pH 7.4) with 0.5% DMSO. Equivalent volumes of labelled protein and compound were mixed. The labelled protein was at a final concentration of 25 nM. The samples were loaded into Monolith premium capillaries (Catalog# MO-K025), and thermophoresis was measured on a Monolith NT.115 equipment (NanoTemper Technologies) with an IR laser power of 40% and an LED intensity of 60%. Each compound was tested in triplicate. The binding curve was calculated from the gradual difference of thermophoresis between the fluorescent molecules of both unbound and bound states, which is plotted as F_norm_ (defined as F_hot_/ F_cold_) against ligand concentration. The binding constants (*K*_d_) were determined from the binding curve by fitting to log(agonist) vs. response in GraphPad Prism version 9.5.1. for Windows, GraphPad Software, San Diego, California USA, www.graphpad.com.

### ChromlogD Assay

The chromlogD assay was carried out on an Agilent 1260 Infinity II^®^ system with a Poroshell 120 EC-C_18_ column [4 µM, 4.6 × 100 mm]; [95:5 H_2_O (50 mM NH_4_.OAc): MeCN → 5:95 H_2_O (50 mM NH_4_.OAc): MeCN, with starting mobile phase at pH 7.4, 10 min; 5 min hold; 1 ml min^−1^]. The Chromatographic Hydrophobicity Index (CHI)^64,78^ values were derived directly from the gradient retention times using calibration parameters for standard compounds. The CHI value approximates to the volume % organic concentration when the compound elutes. CHI was linearly transformed into a chromlogD value by least-square fitting of experimental CHI values using the following formula: chromlogD = 0.0857*CHI - 2^79^.

### Leishmania Promastigote Cell Viability Assay

*L. mexicana* (MNYC/BZ/62/M379) and *L. donovani* LV9 (MHOM/ET/67/HU3) promastigotes were grown at 25 °C in hemoflagellate-modified minimum essential medium (HOMEM) (Gibco) supplemented with 10% (v/v) heat-inactivated foetal calf serum (hi-FCS) (Gibco) and 1% (v/v) penicillin/ streptomycin solution (Sigma-Aldrich). Cell cultures were passaged weekly into fresh medium using dilutions of 1/40 or 1/100 culture in fresh medium. Cell density was determined by fixing cells using a 1/10 dilution in 2% (v/v) formaldehyde and manually counted using a Neubauer haemocytometer.

For cell viability dose-response assays, cultures were grown to mid-log (exponential) phase and dilutions prepared to 5 x 10^3^ cells/ml in medium (HOMEM with 10% hi-FCS and 1% penicillin/ streptomycin solution). 60 µM compound solutions were also prepared in medium. 100 µl cells were seeded into triplicate wells of 96-well plates at 500 cells per well, alongside 100 µl compounds serially diluted in medium to obtain final concentrations of 0 – 30 µM. A miltefosine positive control was included, along with DMSO, parasites-only and media-only controls. Empty wells were filled with 200 µl PBS then plates incubated for 5 days at 25 °C, after which time, 40 µl resazurin (Sigma-Aldrich) added to each well (final concentration 80 µM) and plates incubated for a further 8 hours at 25 °C. Fluorescent readings were taken using a BMG LABTECH CLARIOstar® microplate reader and data processed using BMG LABTECH Mars software with subsequent analysis and graph plotted in GraphPad Prism®. Mean, blank-corrected fluorescence measurements (over protein concentrations 0.23 – 30 µM) were normalised to give values as % cell viability. Three biological replicates were performed, with fluorescence measurements averaged, and data fitted to an [Inhibitor] vs. normalized response -- variable slope dose-response curve to calculate EC_50_. Standard error of the mean was calculated for the averaged mean values.

## Acknowledgements

- C.N.R. thanks the BBSRC for studentship support (BB/M011151/1).
- N.G.J & J.C.M were supported by GSK through the Pipeline Futures Group and a Fellowship from a Research Council United Kingdom Grand Challenges Research Funder under grant agreement ‘A Global Network for Neglected Tropical Diseases’ grant number MR/P027989/1. This work was part-funded by the Wellcome Trust [ref: 204829] through the Centre for Future Health (CFH) at the University of York.
- J.L.C. and S. J. C. thank the EPSRC and GlaxoSmithKline for studentship support (EP/R513295/1). S. J. C. thanks St Hugh’s College, Oxford, for research support.
- We would like to acknowledge our colleagues in The Bioscience Technology Facility of the University of York, including Dr Jared Cartwright and Rebecca Preece for protein production, and Dr Andrew Leech for carrying out SEC-MALLS.
- We are grateful to Sam Hart and Dr Johan Turkenburg for their assistance in X-ray data collection and analysis.
- We thank the DIAMOND Light Source for access to beamline I04 (proposal number mx-18598) that contributed to the results presented here.
- We also thank Dr Raymond Hui and the Structural Genomics Consortium for gifting compounds and plasmids.

## Supplementary

**Table S1.**
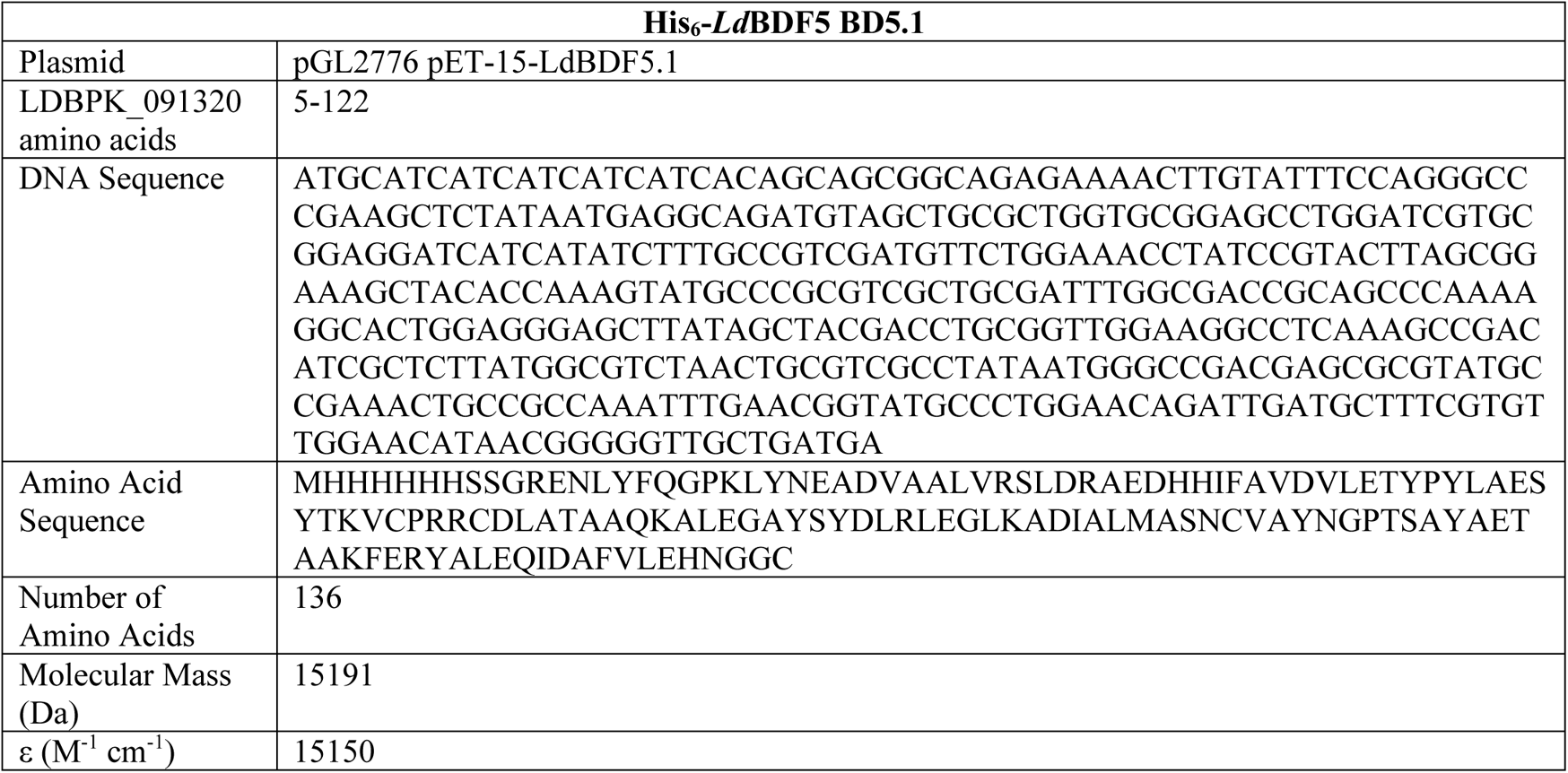
*Ld*BDF5 BD5.1 recombinant protein details; DNA sequence codon-optimised for *E. coli*.

**Table S2.**
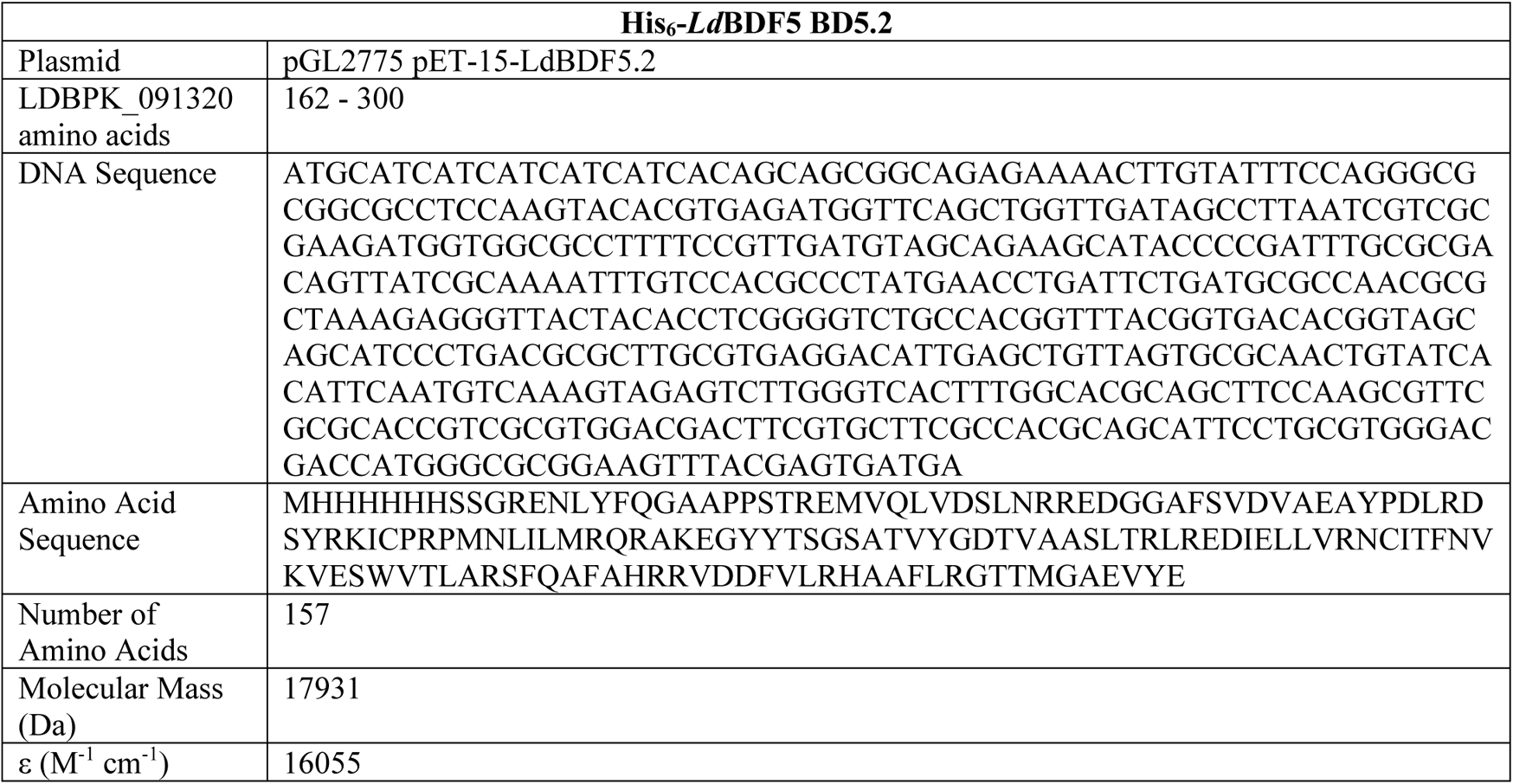
*Ld*BDF5 BD5.2 recombinant protein details; DNA sequence codon-optimised for *E. coli*.

**Table S3.**
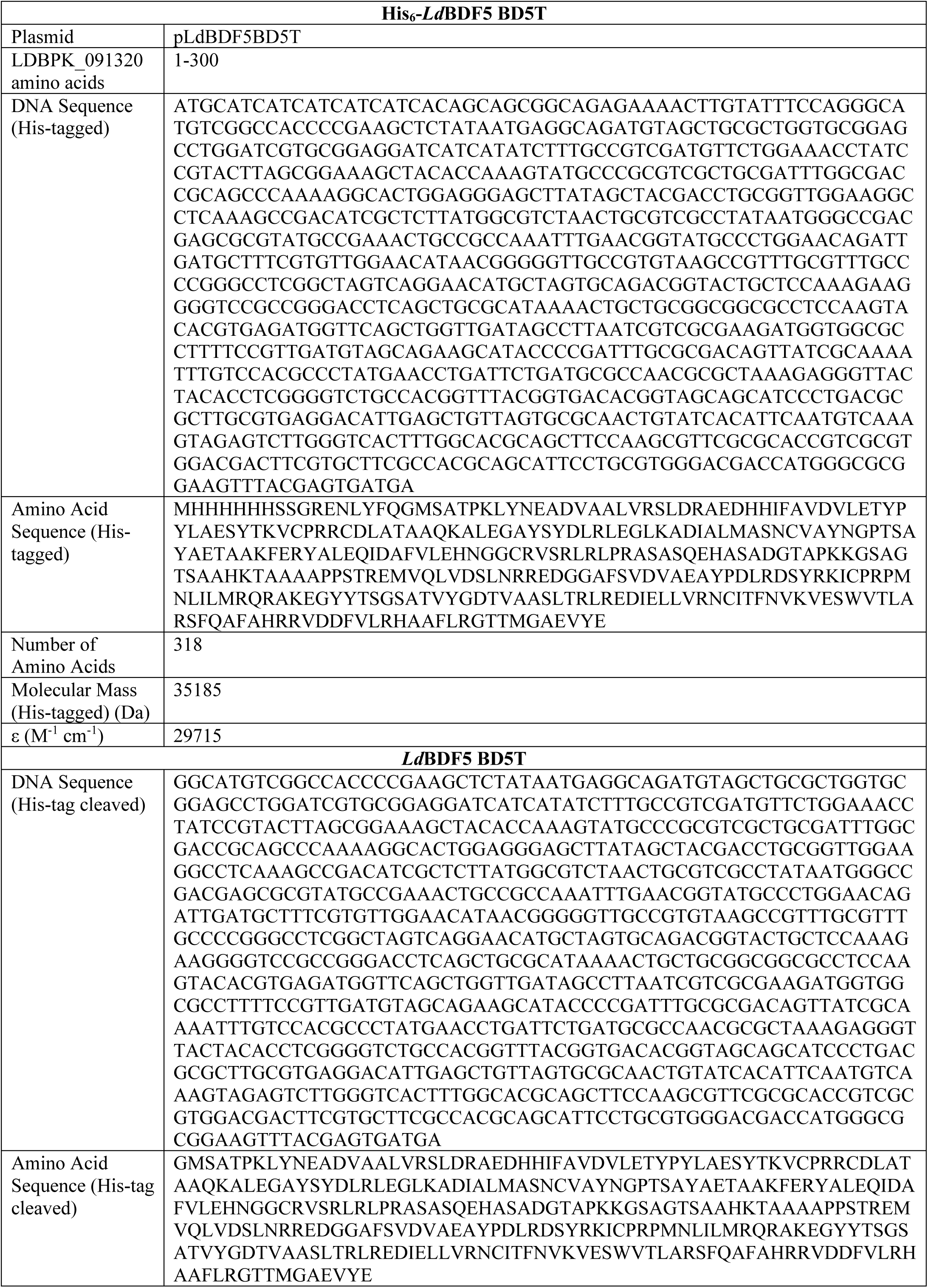
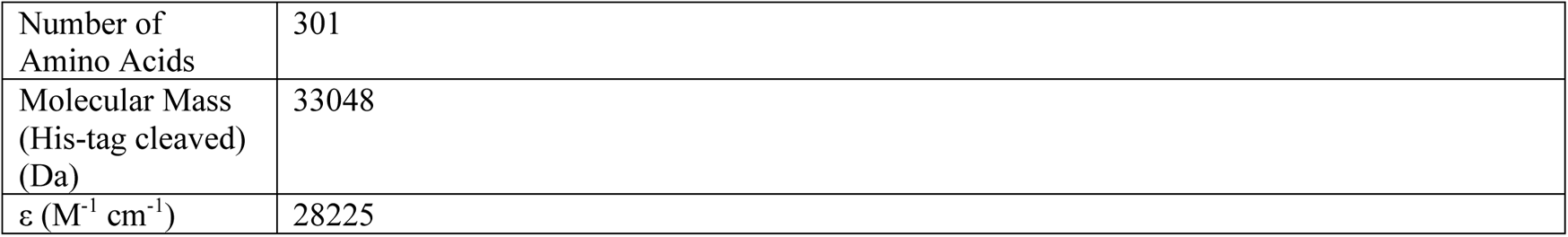
His-tagged and His-tag cleaved *Ld*BDF5 BD5T recombinant protein details; DNA sequence codon-optimised for *E. coli*.

**Table S4.**
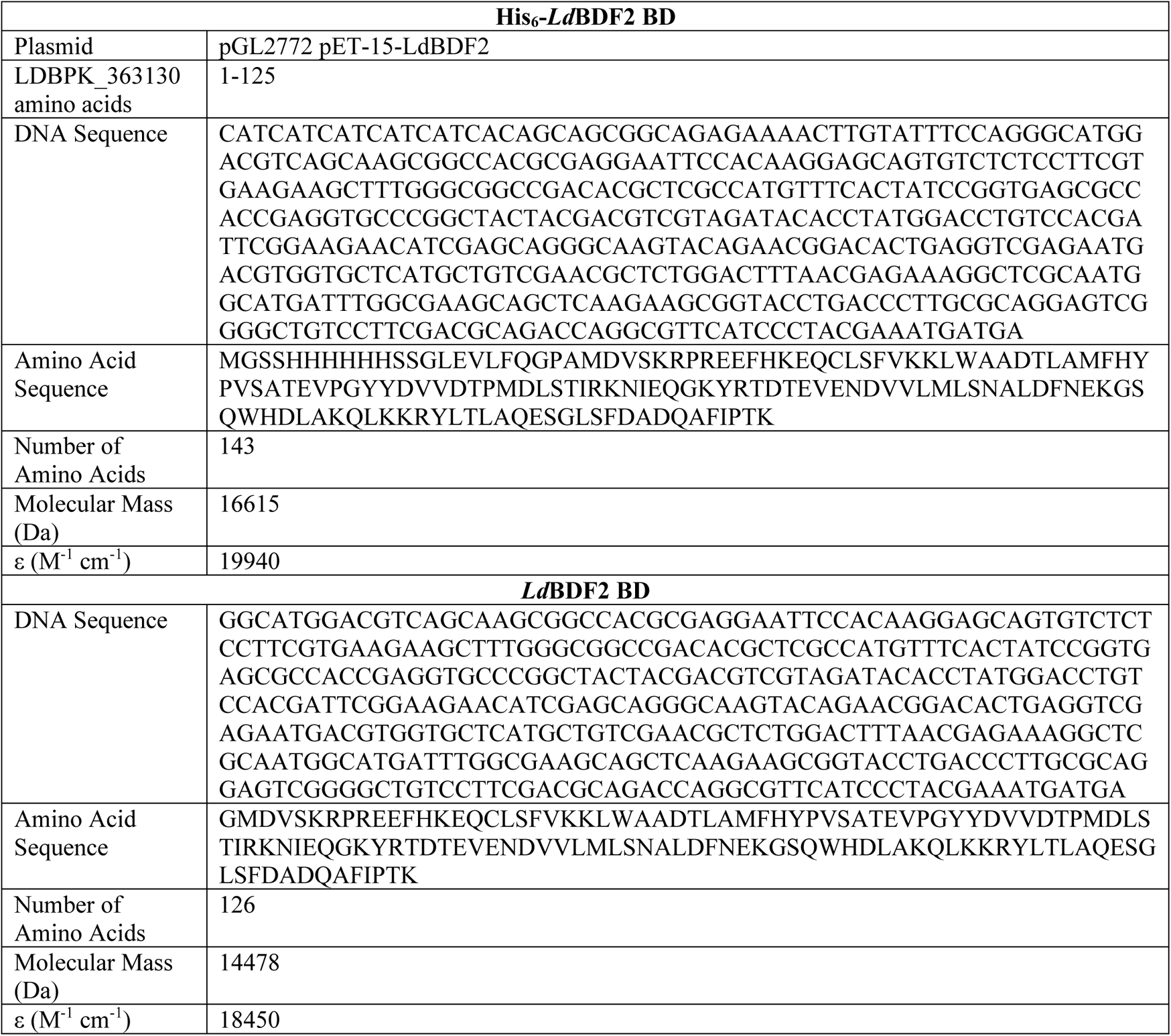
His-tagged and His-tag cleaved *Ld*BDF2 recombinant protein details.

**Table S5.** *Ld*BDF5 BD5.2 apo structure crystallographic data collection and refinement statistics. ^a^ Values in parentheses correspond to the outer resolution shell. *R*_merge_ = ∑*_hkl_*∑*_i_*|*I_i_ −<I> |/*∑*_hkl_*∑*_i_* where *I_i_* is the intensity of the *i*th measurement of a reflection with indexes *hkl* and *<I>* is the statistically weighted average reflection intensity. ^d^ *Rpim* = ∑*_hkl_*[1/(n-1)]^1/2^∑i|Ii - <I>|/∑hkl∑iI *R*work = ∑||*F_o_*| *-* |*F_c_*||*/*∑|*F_o_*| where *F_o_* and *F_c_* are the observed and calculated structure factor amplitudes, respectively. *R*free is the *R*-factor calculated with 5% of the reflections chosen at random and omitted from refinement. Root-mean-square deviation of bond lengths and bond angles from ideal geometry. Percentage of residues in most-favoured/additionally allowed/generously allowed/disallowed regions of the Ramachandran plot, according to PROCHECK.

|  |  |
| --- | --- |
|  | *Ld*BDF5 BD5.2 PDB ID: **8BPT** |
| **Data collection** |  |
| Beamline/Date | i04 01/07/2018 |
| Wavelength (Å) | 0.979500Å |
| Space Group | P2<sub>1</sub>2<sub>1</sub>2<sub>1</sub> |
| Cell dimensions |  |
| *a*, *b*, *c* (Å) | 33.4 75.4 105.8 |
| $\alpha$, $\beta$, $\gamma$ (°) | 90.0, 90.0, 90.0 |
| Resolution (Å) <sup>a</sup> | 31.82 - 1.60 (1.63-1.60) |
| $R_{\text{merge}}$ <sup>a,b</sup> | 0.049 (0.600) |
| $R_{\text{meas}}$ <sup>a,c</sup> | 0.053 (0.639) |
| $R_{\text{pim}}$ <sup>a,d</sup> | 0.018 (0.217) |
| CC(1/2) | 0.999 (0.928) |
| $I / (\sigma I)$ <sup>a</sup> | 19.5 (2.9) |
| Completeness (%) <sup>a</sup> | 98.3 (97.1) |
| Observations <sup>a</sup> | 285,068 (14,599) |
| Unique Reflections <sup>a</sup> | 35,399 (1,738) |
| Wilson B estimate (Å<sup>2</sup>) | 19.20 |
| **Refinement** |  |
| Resolution (Å) | 31.82-1.60 |
| No. reflections | 35,359 (1,735) |
| $R_{\text{work}} / R_{\text{free}}$ | 0.171 / 0.230 |
| No. atoms |  |
| Protein | 2212 |
| Water | 147 |
| *B*-factors |  |
| Protein | 18.01 |
| Water | 32.98 |
| R.m.s. deviations |  |
| Bond lengths (Å) | 0.0138 |
| Bond angles (°) | 1.874 |
| Ramachandran Statistics (%) |  |
| Preferred | 98.91 |
| Allowed | 1.09 |
| Outliers | 0 |

**Figure S1.**
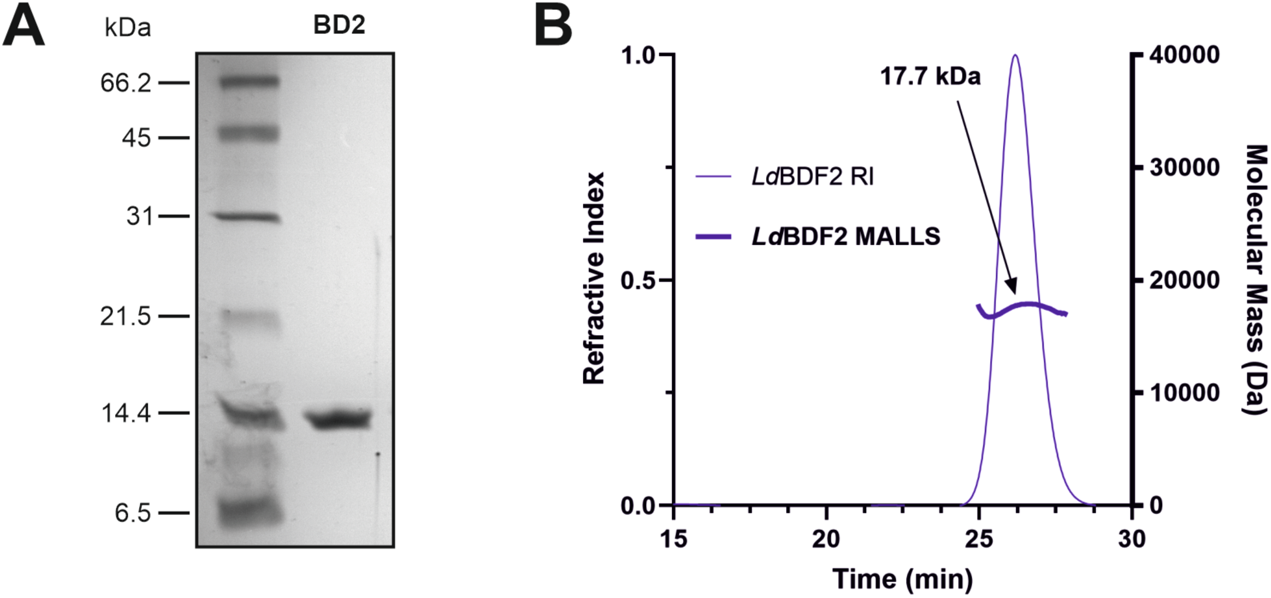
(A) 17.5% SDS-PAGE analysis of purified recombinant His-tag-cleaved BD2 with an expected molecular mass of 14.5 kDa. (B) SEC-MALLS analysis of recombinant His-tagged BD2 with arrow indicating MALLS curve labelled with the associated estimated molecular mass (predicted molecular mass 16.6 kDa).

**Figure S2.**
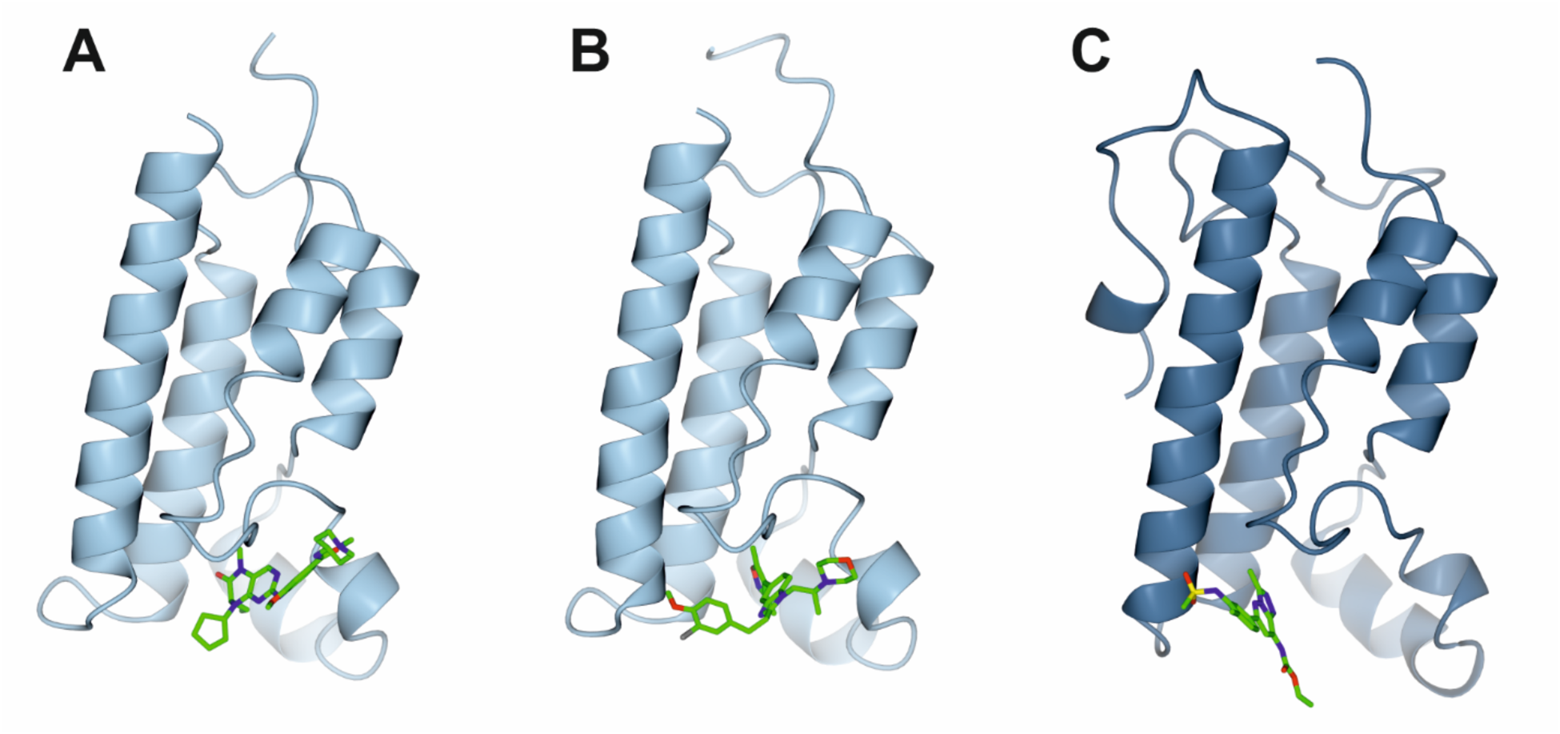
X-ray co-crystal structures of (A) *Ld*BDF BD5.1 in complex with BI 2536, (B) *Ld*BDF BD5.1 in complex with SGC-CBP30, and (C) *Ld*BDF5 BD5.2 in complex with bromosporine. PDB codes are 5TCM, 6BYA & 5TCK, respectively. Figures generated using CCP4mg software.

**Figure S3.**
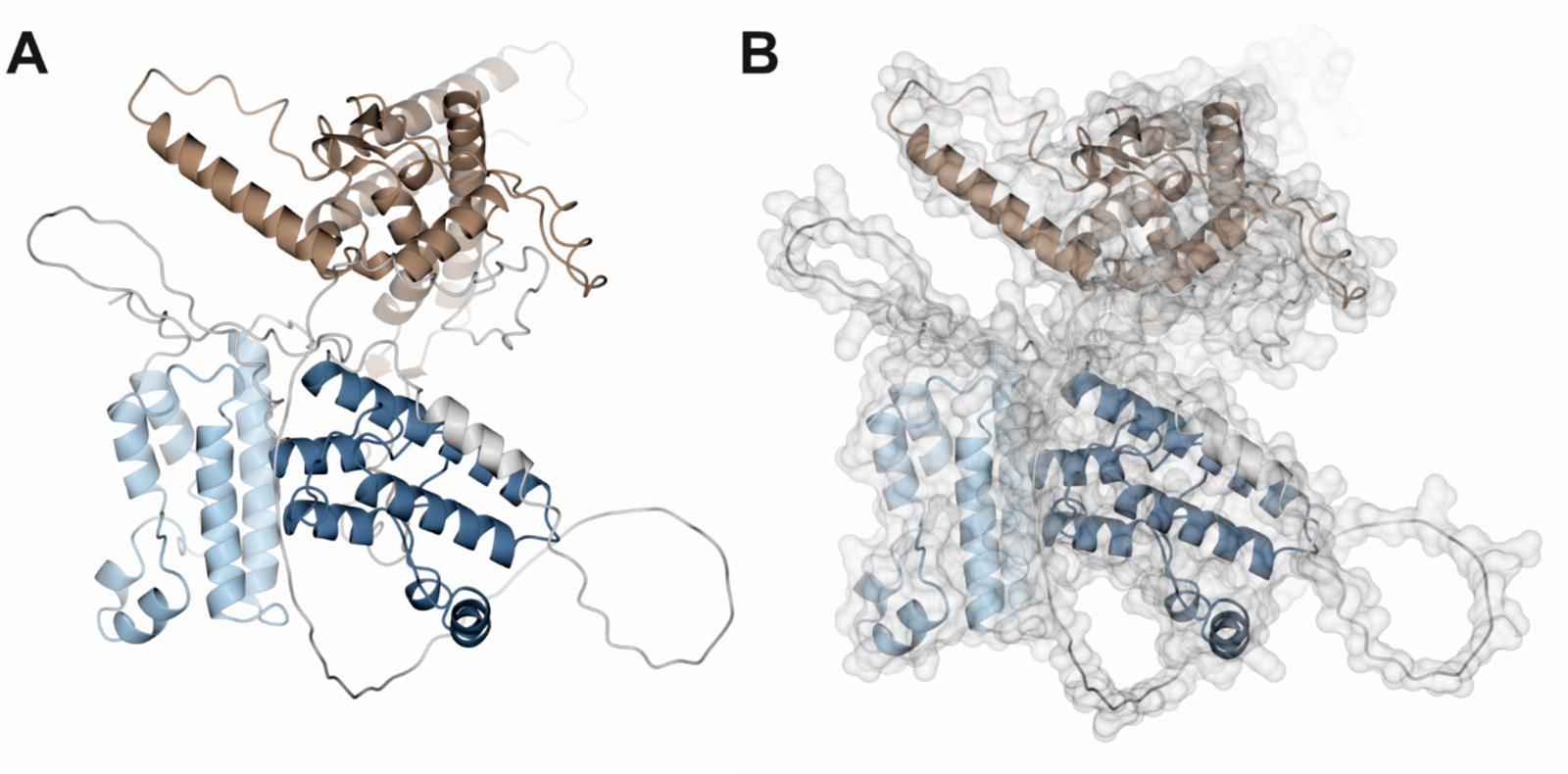
AlphaFold predicted structure of *Ld*BDF5 (LINF_090020000) showing (A) ribbon and (B) surface representations. Different colours indicate the different domains, BD5.1 (light blue), BD5.2 (dark blue) and predicted C-terminal MRG domain (brown).

**Figure S4.**
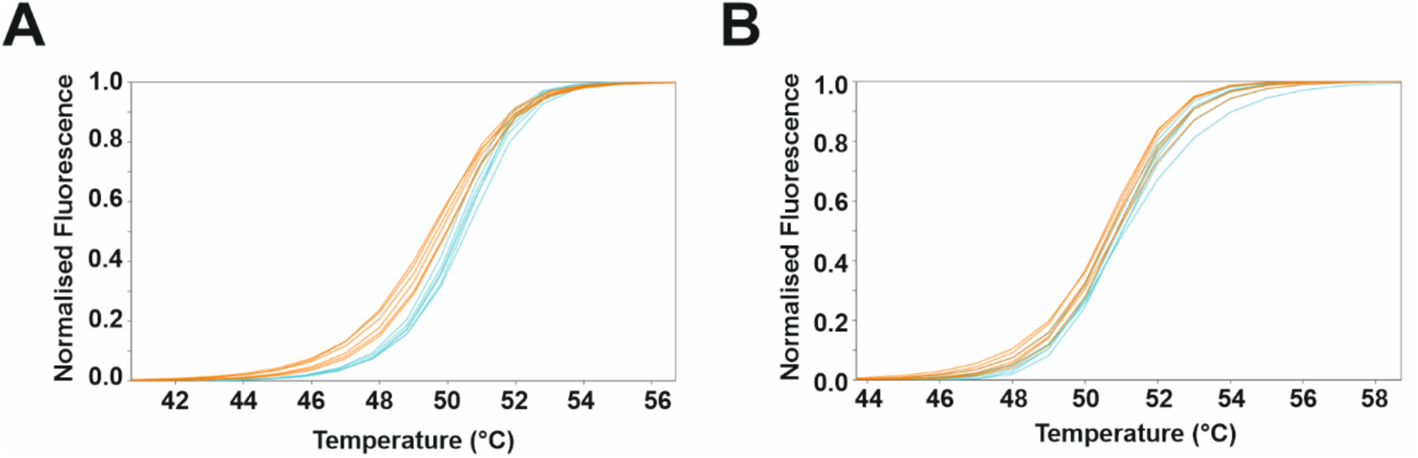
Thermal shift assay melting curves for *Ld*BDF5 BD5T with (A) pan-acetylated H2B_9-23_K9^Ac^K15^Ac^K19^Ac^K21^Ac^ and (B) unmodified H2B_9-23_ peptides at 400 µM. Curves are normalised for five parameter sigmoid equation model fitting for six replicate samples of protein with peptides (blue) alongside DMSO control samples (orange). Graphs produced using the online JTSA tool.

**Figure S5.**
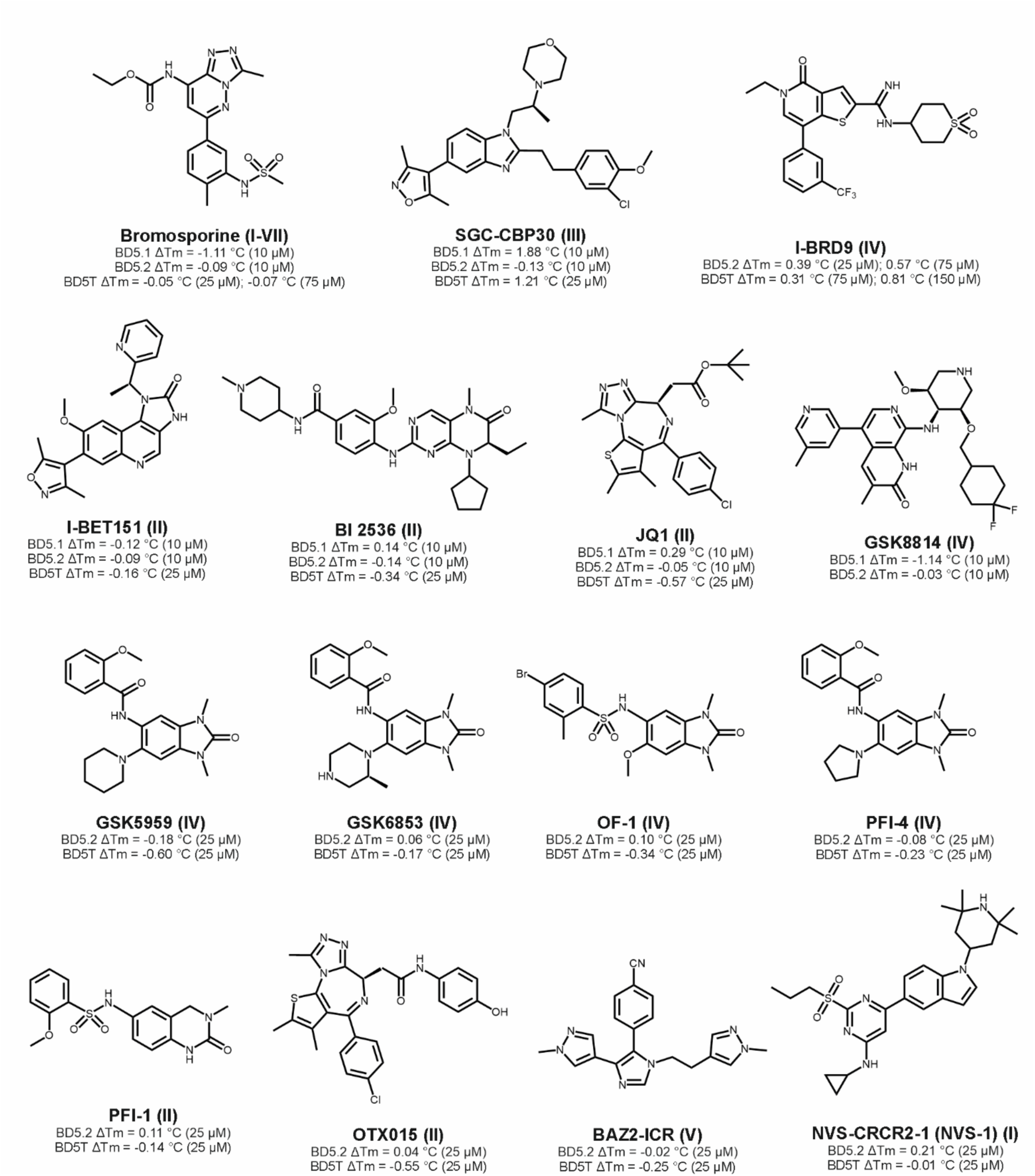
Structures of 15 human bromodomain inhibitor compounds and the family of human protein target(s) in brackets. Thermal shifts from TSA screen with the *Ld*BDF5 recombinant proteins given below, where compound concentrations are given in brackets.

**Figure S6.**
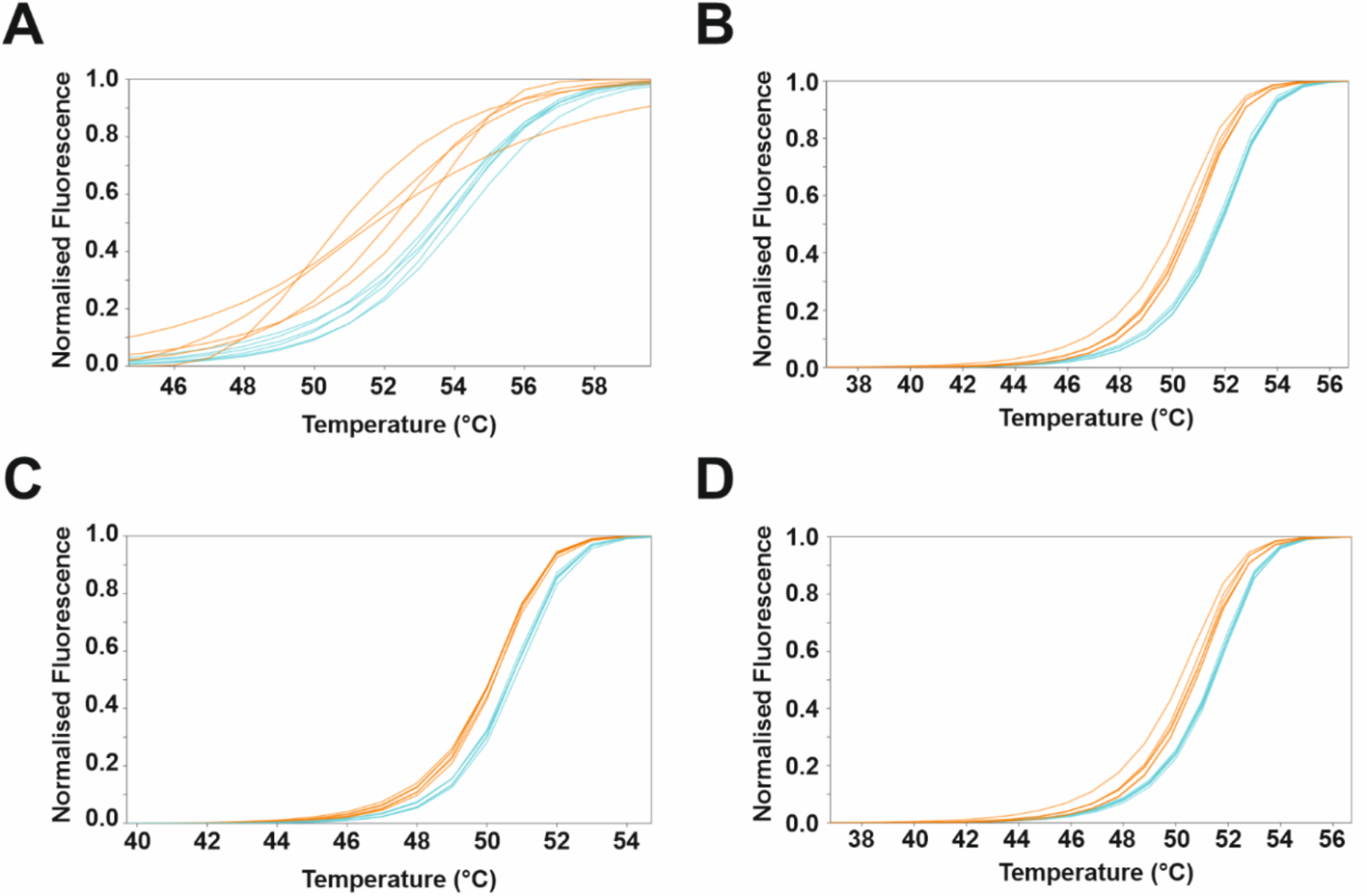
Thermal shift assay melting curves for (A) BD5.1 + SGC-CBP30 at 10 µM, (B) BD5T + SGC-CBP30 at 25 µM, (C) BD5.2 + I-BRD9 at 75 µM, and (D) BD5T + I-BRD9 at 150 µM. Curves are normalised for five parameter sigmoid equation model fitting for five or six replicate samples of protein with compounds (blue) alongside DMSO control samples (orange).

**Figure S7.**
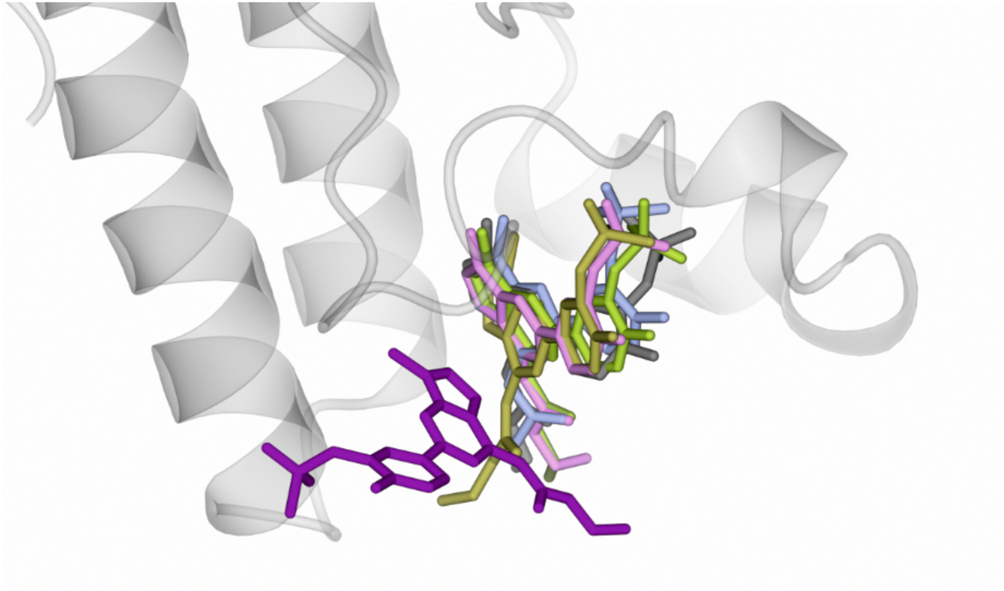
Overlay of co-crystal structures showing bromosporine binding to bromodomains of *Ld*BDF5 BD5.2 (PDB code 5TCK, purple); *Ld*BDF2 (PDB code 5C4Q, blue); *Ld*BDF3 (PDB code 5FEA, yellow), human BRD4 (PDB code 5IGK, grey); human BRD7 (PDB code 6V1H, pink); and human BRD9 (PDB code 5IGM, green). Protein structure of *Ld*BDF5 BD5.2 is shown as ribbon representation.

**Figure S8.**
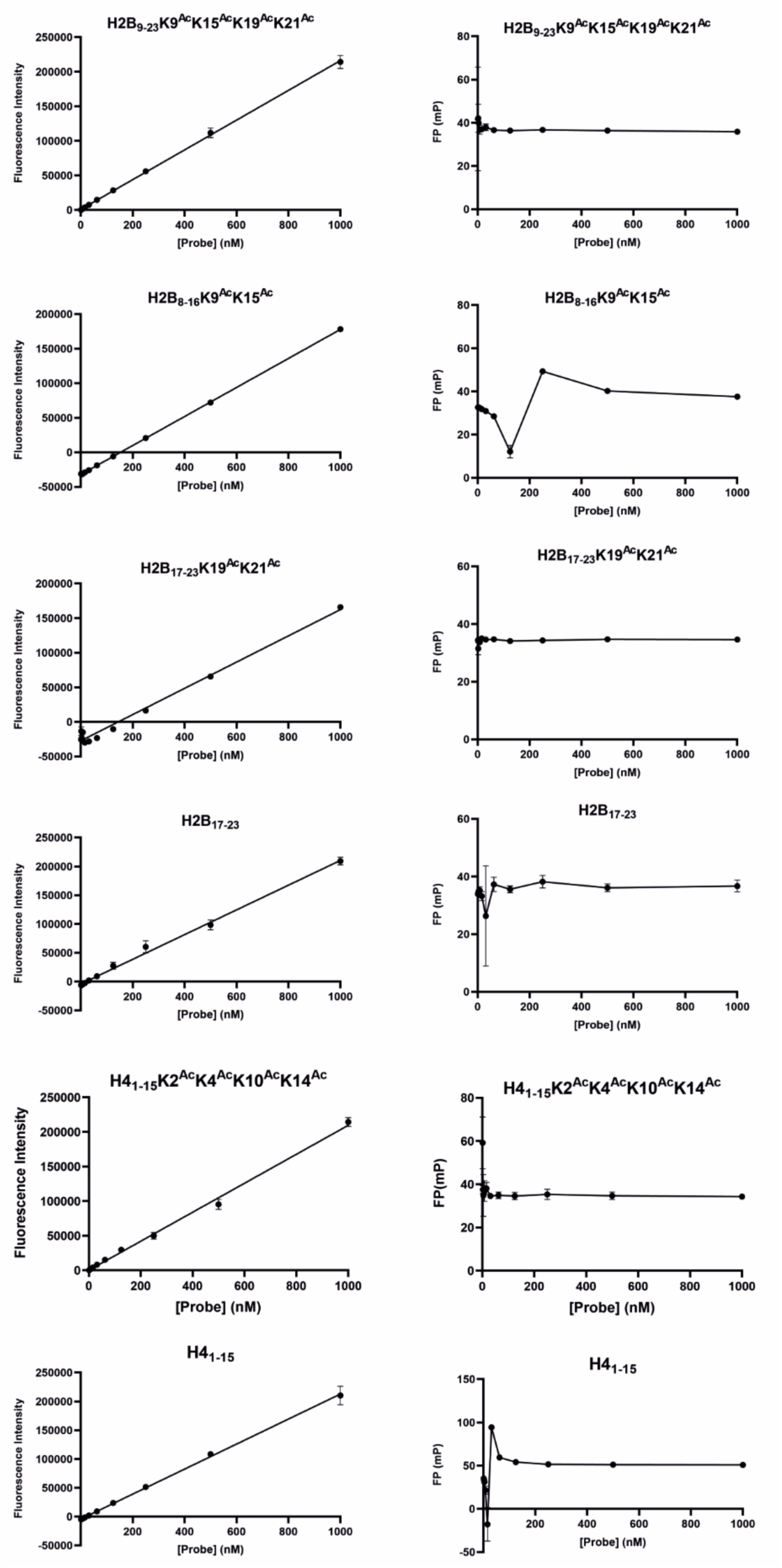
FP probe optimisation with (A) fluorescence intensity and (B) fluorescence polarisation recorded for each peptide probe. Mean, blank-corrected values are plotted against probe concentration and fluorescence intensity data is fitted to linear regression. Error bars represent SD (n = 3).

## Notes

### Competing Interest Statement

We disclose that Felix Calderon, Raquel Gabarro and Jacob Bush are employees of GSK. GSK provided some of the SGC Tool Compounds to support the lab work. This work was in part supported by funding from GSK through the Pipeline Futures Group.

